# Sensory feedback independent pre-song vocalizations correlate with current state of motor preparation

**DOI:** 10.1101/509505

**Authors:** Divya Rao, Satoshi Kojima, Raghav Rajan

**Author notes:** Correspondence should be addressed to Raghav Rajan.

## Abstract

Many self-initiated, learned, motor sequences begin by repeating a simple movement, like ball-bouncing before a tennis serve, and this repetition is thought to represent motor preparation. Do these simple movements provide real-time sensory feedback used by the brain for getting ready or do they simply reflect internal neural preparatory processes? Here, we addressed this question by examining the introductory notes (INs) that zebra finches repeat before starting their learned song sequence. INs progress from a variable initial state to a stereotyped final state before each song and are thought to represent motor preparation before song. Here, we found that the mean number of INs before song and the progression of INs to song were not affected by removal of two sensory feedback pathways (auditory and proprioceptive). In both feedback-intact and feedback-deprived birds, the presence of calls (other non-song vocalizations), just before the first IN, was correlated with fewer INs before song and an initial state closer to song. Finally, the initial IN state correlated with the time to song initiation. Overall, these results show that INs do not provide real-time sensory feedback for preparing the motor system. Rather, repetition of INs, and possibly, other such simple movements, may reflect the “current” state of internal neural preparatory processes involved in getting the brain ready to initiate a learned movement sequence.

**SUMMARY STATEMENT:** The number and progression of introductory notes to song in the zebra finch are not affected by removal of sensory feedback.

## INTRODUCTION

All movements are believed to be planned in the brain before execution (Shenoy et al., 2011; Shenoy et al., 2013; Svoboda and Li, 2018; Wong et al., 2015). Such planning is characterized by changes in neural activity seconds to hundreds of milliseconds before the start of the movement. Such changes in activity have been observed in many different brain areas in humans (Fried et al., 2011) and in other model organisms (Chen et al., 2017; Churchland et al., 2006a; Churchland et al., 2006b; Gao et al., 2018; Lee and Assad, 2003; Li et al., 2015; Maimon and Assad, 2006; Murakami and Mainen, 2015; Murakami et al., 2014; Romo and Schultz, 1987; Tanji and Evarts, 1976). While all of these studies have focused on simple movements, a number of human and animal movements consist of learned sequences of simple movements – for eg. the serve of a tennis player or the song of a bird. Such movement sequences are often preceded by the repetition of a simple movement, like the repeated bouncing of a ball before the tennis serve, and this repetition of a simple movement is thought to represent motor preparation (Cotterill, 2010). How do these simple movements help prepare the brain for the execution of a more complex motor sequence? Do they provide sensory feedback that is used by the brain for preparation or do these simple movements simply represent internal neural preparatory processes? Here, we addressed this question using the song of the adult male zebra finch as our model system.

The song motif (referred to as song) of the adult male zebra finch, consisting of a stereotyped sequence of sounds (syllables) interleaved with silent gaps (Fig 1A), is a well-established model for understanding movement sequences (Fee and Scharff, 2010). Song is learned by young birds from a conspecific tutor during a critical period (Fee and Scharff, 2010). While song is typically part of a courtship ritual for mate attraction, birds also sing when they are alone (undirected song) (Sossinka and Böhner, 1980; Zann, 1996), making this an excellent model system to study motor preparation before self-initiated, learned, movement sequences.

**Figure 1.**
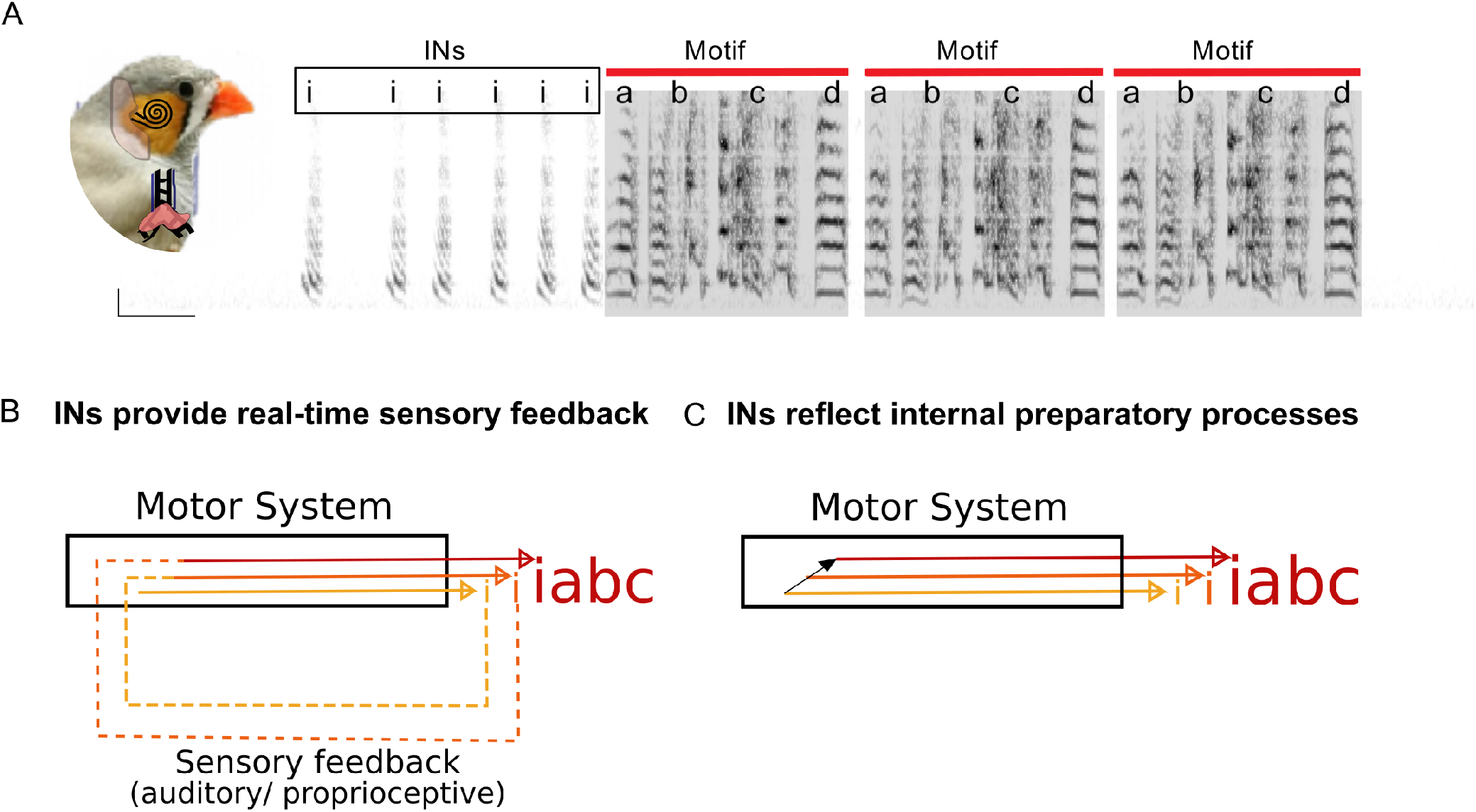
Zebra finch song and possible hypothesis about the role of INs in song initiation. (A) A male zebra finch and spectrogram of the song of an adult male zebra finch. The picture also shows two important sensory feedback sources, namely, auditory feedback from the cochlea and proprioceptive feedback from the syringeal muscles. In the spectrogram, ‘i’ denotes introductory notes (INs) and ‘a’, ‘b’, ‘c’ and ‘d’ represent the syllables of the song motif (gray shading). (B),(C) Hypotheses for the role of INs in song initiation. ‘i’ represents INs and the colour, size and distance between INs represent the changing acoustic and timing features of successive INs. Each IN provides sensory feedback to the brain to drive the next improved IN until song initiation (B) OR Each improved IN reflects changes in ongoing internal preparatory activity (C).

Song, like other movement sequences, is preceded by the bird repeating a short vocalization called an Introductory Note (IN – ‘i’s in Fig. 1A) (Price, 1979; Sossinka and Böhner, 1980). Each song bout consists of a variable number of such INs followed by multiple repeats of the song. We have previously shown that intervals between successive INs and the acoustic properties of successive INs progress from a variable initial state (first IN in each song bout) to a more consistent “ready” state (last IN in each song bout) just before the start of each song (Rajan and Doupe, 2013). Given the similarity to the reduction in variability associated with neural preparatory activity before the onset of simple movements (Churchland et al., 2006c), INs may represent vocalizations that help prepare the zebra finch brain to produce song. However, what INs represent and what role they play in song initiation remains unclear.

To understand what INs represent, it is important to first determine whether each IN provides sensory feedback that is used by the brain to produce an “improved” next IN that is closer to the “ready” state (Fig. 1B). Sensory feedback could also induce adaptation that drives the system forward to the song as has been proposed for the transition from repeating syllables within Bengalese finch song (Wittenbach et al., 2015). Consistent with this hypothesis, previous studies disrupting proprioceptive feedback or auditory feedback have reported changes in the number of INs before song in some birds (Bottjer and Arnold, 1984). However, these changes have not been quantified rigorously and the specificity of these changes to removal of feedback has not been determined.

In contrast to both these hypotheses that require sensory feedback, it is also possible that INs and the progression of INs to song reflect a passive readout of internal preparatory processes (Fig. 1C). Internal preparatory activity has been shown to occur in a number of song control nuclei in the zebra finch brain, hundreds of milliseconds before the start of the first IN of an undirected song bout (Hessler and Doupe, 1999; Kao et al., 2008; Rajan, 2018; Woolley et al., 2014). Further, premotor activity is also present, in premotor and motor nuclei, before the start of each IN (Danish et al., 2017; Rajan and Doupe, 2013; Vyssotski et al., 2016; Yu and Margoliash, 1996). In premotor nucleus HVC, IN-related neural activity becomes distinct before the last IN acting as an early indicator that song is about to start (Rajan and Doupe, 2013). This IN-related activity could also represent a continuation of internal preparatory activity, albeit, associated with overt preparatory vocalizations (Fig. 1C).

To distinguish between these two possibilities (Fig. 1B and Fig. 1C), we analyzed the number and properties of INs soon after removal of two important forms of sensory feedback, namely proprioceptive feedback from the syringeal muscles (Bottjer and Arnold, 1984; Vicario, 1991; Williams and McKibben, 1992) or auditory feedback (Konishi, 1965). We found that mean IN number before song and progression of INs to song were not affected by removal of either form of feedback. Further, the progression of INs to song was not affected by removal of neural input to the syringeal muscles. Finally, we found fewer INs and a quicker transition to song when the first IN was produced soon after calls (non-song vocalizations that are different from INs and song syllables). These data demonstrate that INs do not provide sensory feedback. Rather, INs may reflect the “current” state of ongoing internal neural preparatory processes involved in getting the zebra finch brain “ready” to produce the learned song sequence.

## MATERIALS AND METHODS

Experimental procedures performed at IISER Pune were approved by the Institute Animal Ethical Committee in accordance with the guidelines of the Committee for the Purpose of Control and Supervision of Experiments on Animals (CPCSEA, New Delhi). Experiments performed at UCSF, San Francisco were approved by the UCSF Institutional Animal Care and Use Committee in accordance with NIH guidelines.

### Birds and song recording

All birds (n=42) used in this study were > 100 days post hatch at the time of the experiment and were either purchased from an outside vendor (n=13), bred at IISER Pune (n=17) or bred at UCSF (n=12). Birds were kept in separate sound isolation boxes (Newtech Engineering Systems, Bangalore, India or Acoustic Systems, Austin, TX) for the duration of the experiment. All songs were recorded by placing a microphone (AKG Acoustics C417PP omnidirectional condenser microphone or B3 lavalier microphone, Countryman Associates, CA) at the top of the cage. For the ts-cut and sham-surgery birds, we kept the position of the microphone the same for recording songs before and after surgical manipulations. Signals from the microphone were amplified using a mixer (Behringer XENYX 802) and then digitized on a computer at a sampling rate of 44100 Hz using custom software. Songs were recorded in “triggered” mode before and after surgery, where data was saved when the microphone signal crossed a preset threshold. Along with the data that crossed the threshold, 1-3 s of data before and after threshold crossing were also saved. For a subset of birds, data was saved in “continuous” mode where all of the data for the entire recording period was saved. All songs were recorded in the “undirected” condition. Songs of 3 of the birds used for the analysis of calls and their influence on song initiation have been used in a previous study for analysis of INs before song (Rajan and Doupe, 2013). The influence of calls on song initiation was not considered in the previous study. For the analysis of day-to-day changes in IN number and properties, we used data from 14 birds that were recorded on two different days (range: 1-3 days apart). Of these 14 birds, 1 bird was used at a later time-point for ts-cut surgery with a new set of pre and post surgery recordings and 10 birds were used for analysis of the influence of calls on INs. Pre-surgery recordings for 18/21 birds (n=5 ts-cut, n=6 sham surgery and n=7 deaf) were done 0-2 days before surgery. For the remaining 3 birds (n=3 ts-cut), pre-surgery recordings were done 18, 15 and 5 days before surgery respectively.

### ts nerve cut and sham surgery

Tracheosyringeal nerves were surgically cut using previously described protocols (Bottjer and Arnold, 1984; Vicario, 1991; Williams and McKibben, 1992). Briefly, birds (n=9) were deeply anesthetized by intramuscular injection of ketamine (30mg/kg), xylazine (3 mg/kg) and diazepam (7 mg/kg). Absence of a response to toe pinch was used to assess depth of anesthesia. Birds were then placed on a platform with the ventral side facing up. A rolled tissue under the neck served to stretch and give easy access to the throat. Feathers were plucked and an incision ~10 mm was made. The trachea was exposed by removal of fat tissue. Using fine forceps the tracheosyringeal (ts) nerve bundle on either side of the trachea was pulled away from the trachea and part of the nerve (n=9 birds, median length cut = 4mm; range – 2mm to 7mm) was cut out on both sides using spring scissors (Fine Science Tools, CA, USA). The skin was then glued using tissue adhesive (Vetbond, 3M). For sham surgeries (n=6), the same procedure was followed but the ts nerves were not cut. In two of the sham surgery birds, some cuts were made on the thick membrane enclosing the oesophagus. Birds typically resumed singing within 10 days of surgery. We considered songs produced on the second day of singing after surgery (sham: 2-5 days and ts-cut: 3-10 days after surgery) for analysis because of higher number of songs. For one bird, we did not have pre-surgery songs in the undirected condition, so we excluded this bird for analyses involving comparison to pre-surgery (Fig. 2, 3, 4, 5, 6). Data from this bird was included only for analysis of influence of calls on the number and properties of INs (Fig. 7, 8).

**Figure 2.**
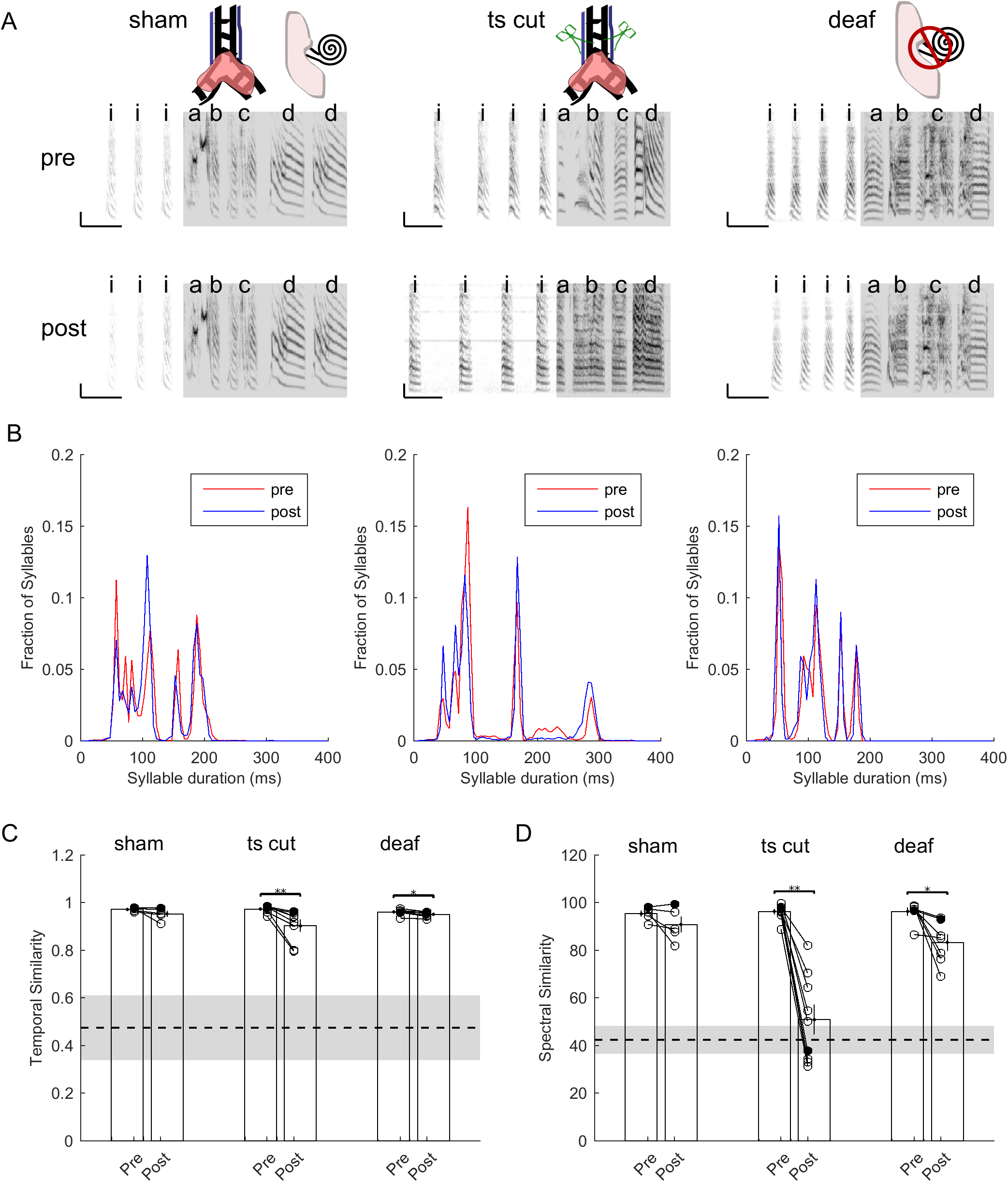
Song structure after sham surgery, tracheosyringeal nerve cut or deafening. (A) Representative spectrograms of song for individual birds before and after sham surgery (left), tracheosyringeal nerve cut (middle) or deafening (right). Scale bar 200ms and 1000Hz. ‘i’s represent INs and ‘a’, ‘b’, ‘c’ and ‘d’ represent song motif syllables. (B) Syllable duration histograms for individual birds before (red) and after (blue) sham surgery (left), ts-cut (middle) or deafening (right). (C) and (D) Temporal similarity (C) and spectral similarity (D) to pre surgery song for songs produced after sham surgery, ts-cut or deafening. Each circle represents one bird and lines connect data from the same bird before and after surgery. Bars and whiskers represent means across birds. Dashed line and shading represents mean and 95% confidence intervals for similarity between random birds. Filled circles represent values for the birds shown in (A) and (B).

**Figure 3.**
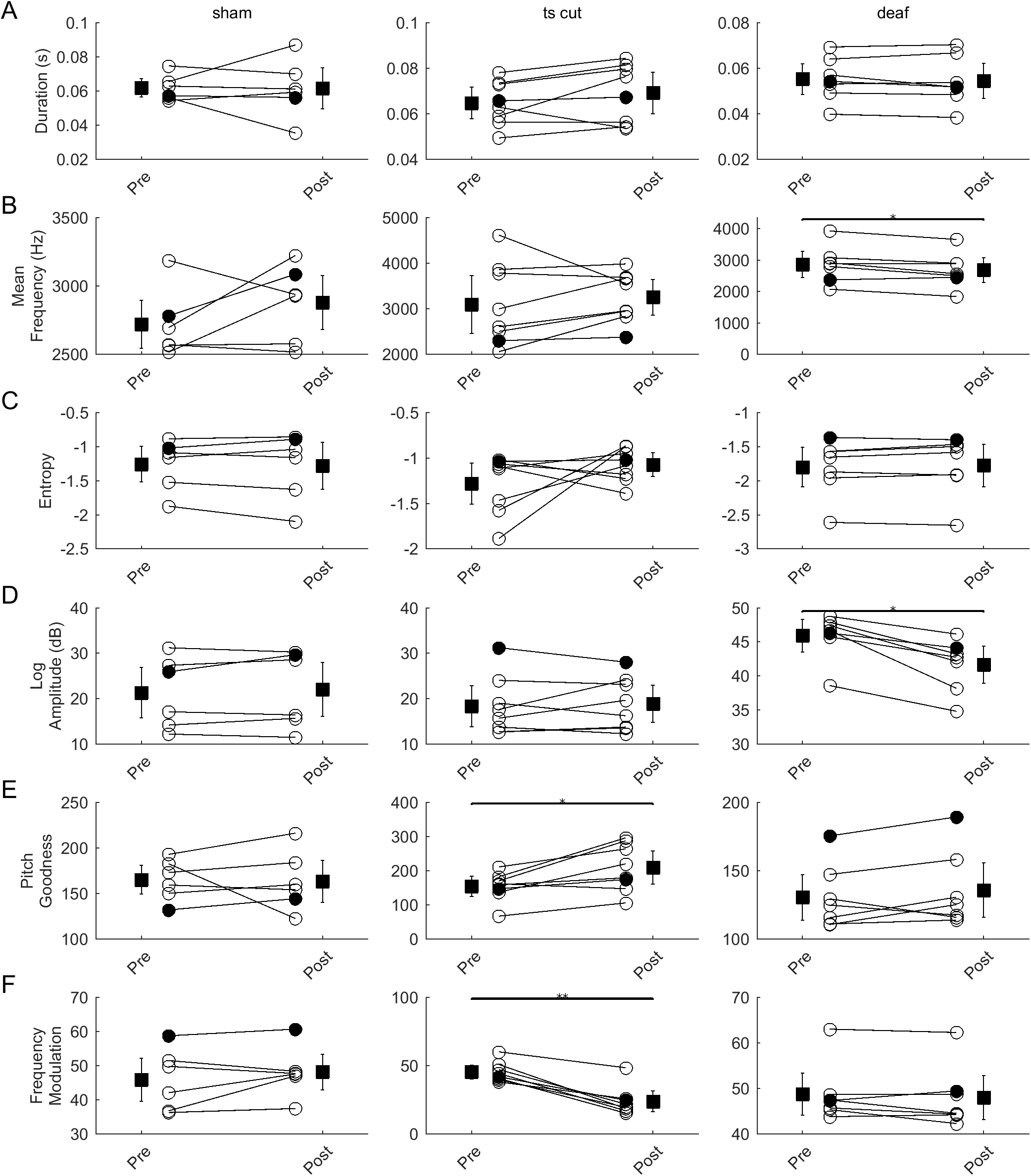
Changes in IN acoustic features after removal of sensory feedback. (A), (B), (C), (D), (E) and (F) Acoustic properties of INs pre and post sham surgery (left column), ts-cut surgery (middle column) or deafening (right column). Circles represent individual birds and lines connect data from the same bird. Squares and whiskers represent mean and s.e.m. across birds. Acoustic features plotted are duration (A), mean frequency (B), entropy (C), log amplitude (D), pitch goodness (E) and frequency modulation (F). * represents p < 0.05, ** represents p < 0.01, Wilcoxon sign-rank test.

**Figure 4.**
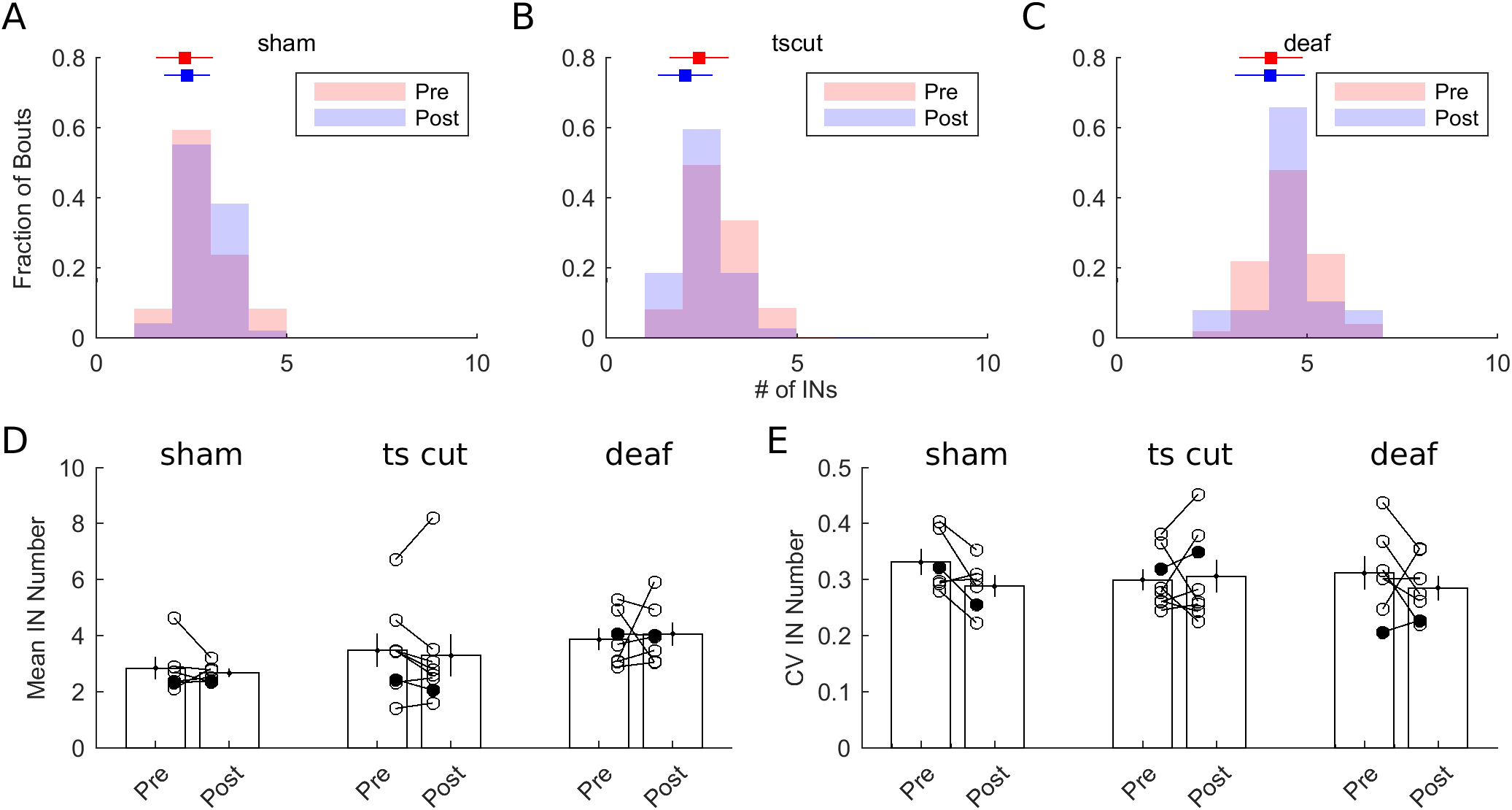
Mean IN number and variability are not affected by removal of sensory feedback. (A), (B) and (C) Distribution of number of INs for a representative bird before (red) and after (blue) sham-surgery (A), ts-cut (B) and deafening (C). Squares and whiskers represent mean and s.d. of the distributions pre (red) and post (blue) surgery. (D) and (E) Mean IN number (D) and CV of IN number (E) before and after surgery for all sham-surgery, ts-cut and deaf birds. Circles represent individual birds and lines connect data from the same bird. Bars and whiskers represent mean and s.e.m. across birds. Filled circles represent the birds shown in (A), (B) and (C)

**Figure 5.**
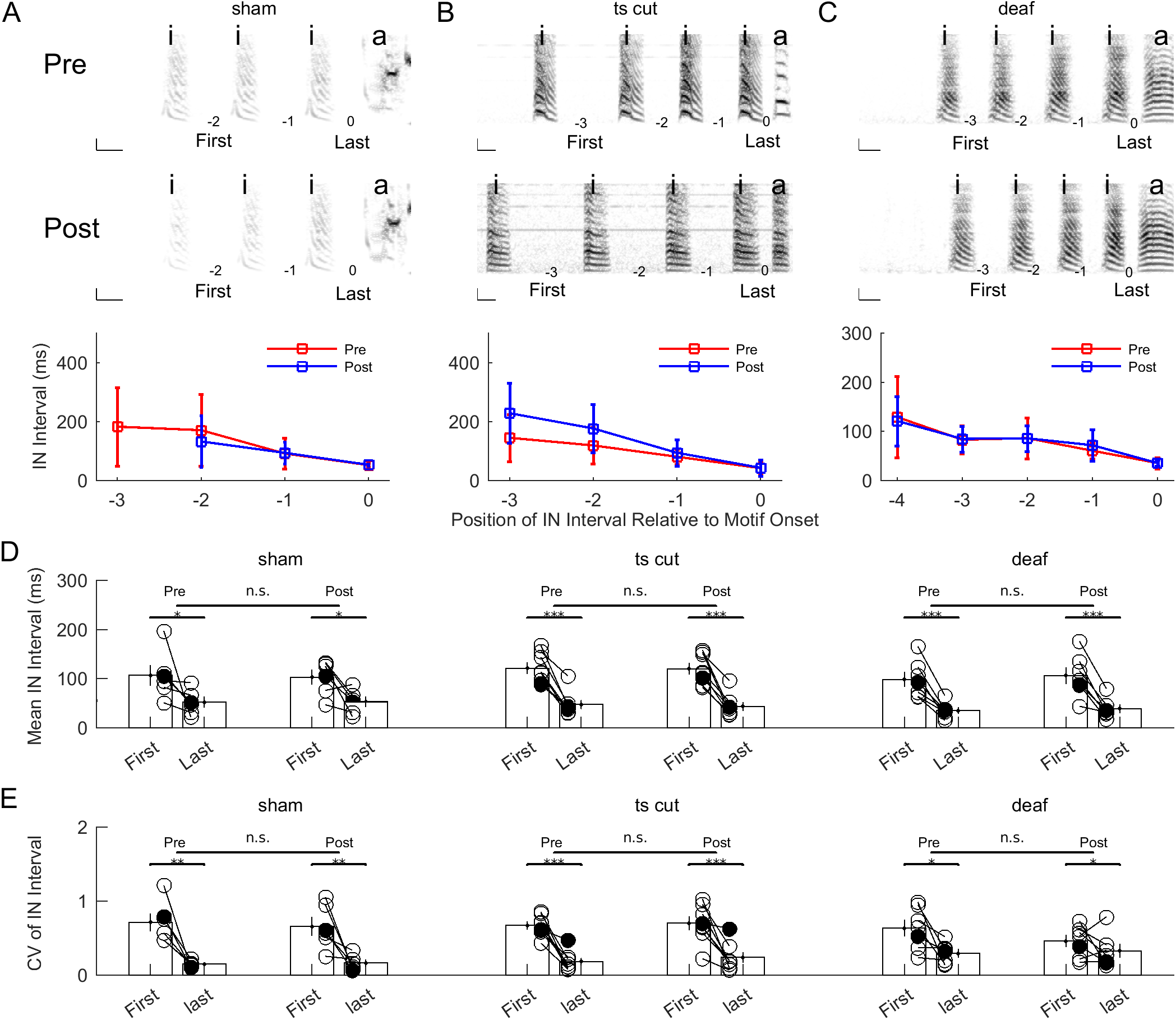
Progression of intervals between successive INs is independent of sensory feedback. (A),(B) and (C) Top – spectrograms of a sequence of INs before the first motif syllable before and after surgery. The position of each IN interval relative to the motif onset and the first and last interval are marked. Bottom – Interval between successive INs vs position of the interval relative to motif onset for 3 example birds before (red) and after (blue) sham surgery (A), ts-cut surgery (B) or deafening (C). Squares and whiskers represent mean and standard deviation. (D) Mean interval between the first two INs in a bout and mean interval between the last IN and song across all birds before and after sham surgery (left), ts-cut surgery (middle) or deafening (right). Circles represent individual birds and lines connect data from the same bird before and after surgery. Bars and whiskers represent mean and s.e.m. across all birds. (E) CV (standard deviation / mean) of the interval between the first two INs in a bout and CV of the interval between the last IN and song across all birds before and after sham surgery (left), ts-cut surgery (middle) or deafening (right). Circles represent individual birds and lines connect data from the same bird before and after surgery. Bars and whiskers represent mean and s.e.m. across all birds. * represents p < 0.05, ** represents p < 0.01, *** represents p < 0.001, Repeated Measures 2-way ANOVA.

**Figure 6.**
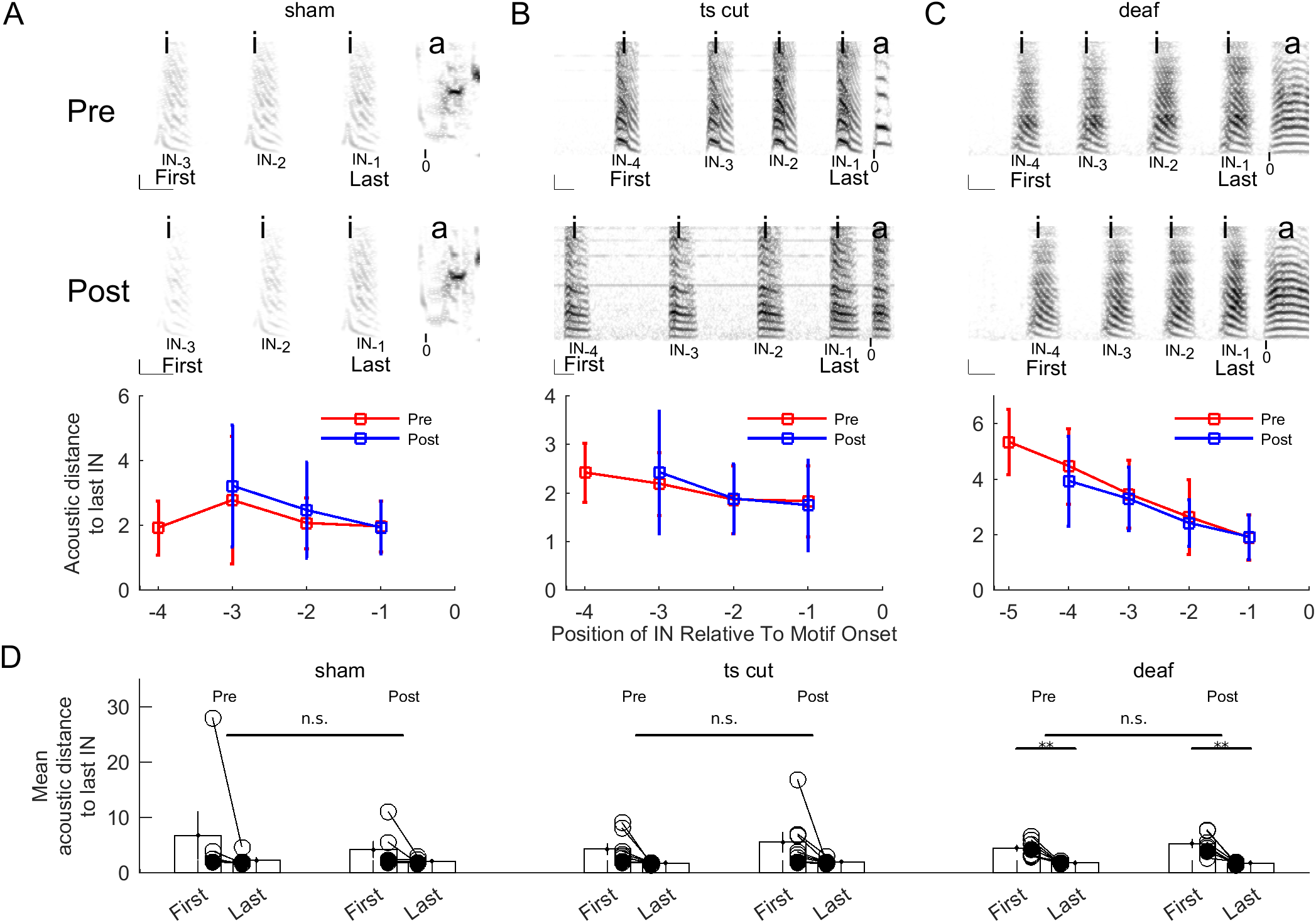
Progression of acoustic properties of successive INs is independent of sensory feedback. (A), (B) and (C) Top – spectrograms of a sequence of INs before the first motif syllable before and after surgery. The position of each IN relative to motif onset and the first and last INs are marked. Bottom – Acoustic distance of successive INs from the last IN for 3 example birds before (red) and after (blue) sham surgery (A), ts-cut surgery (B) or deafening (C). Squares and whiskers represent mean and standard deviation. (D) Mean acoustic distance of the first IN and the last IN from the last IN in a bout across all birds before and after sham surgery (left), ts-cut surgery (middle) or deafening (right). Circles represent individual birds and lines connect data from the same bird before and after surgery. Bars and whiskers represent mean and s.e.m. across all birds. * represents p < 0.05, ** represents p < 0.01, *** represents p < 0.001, Repeated Measures 2-way ANOVA.

**Figure 7.**
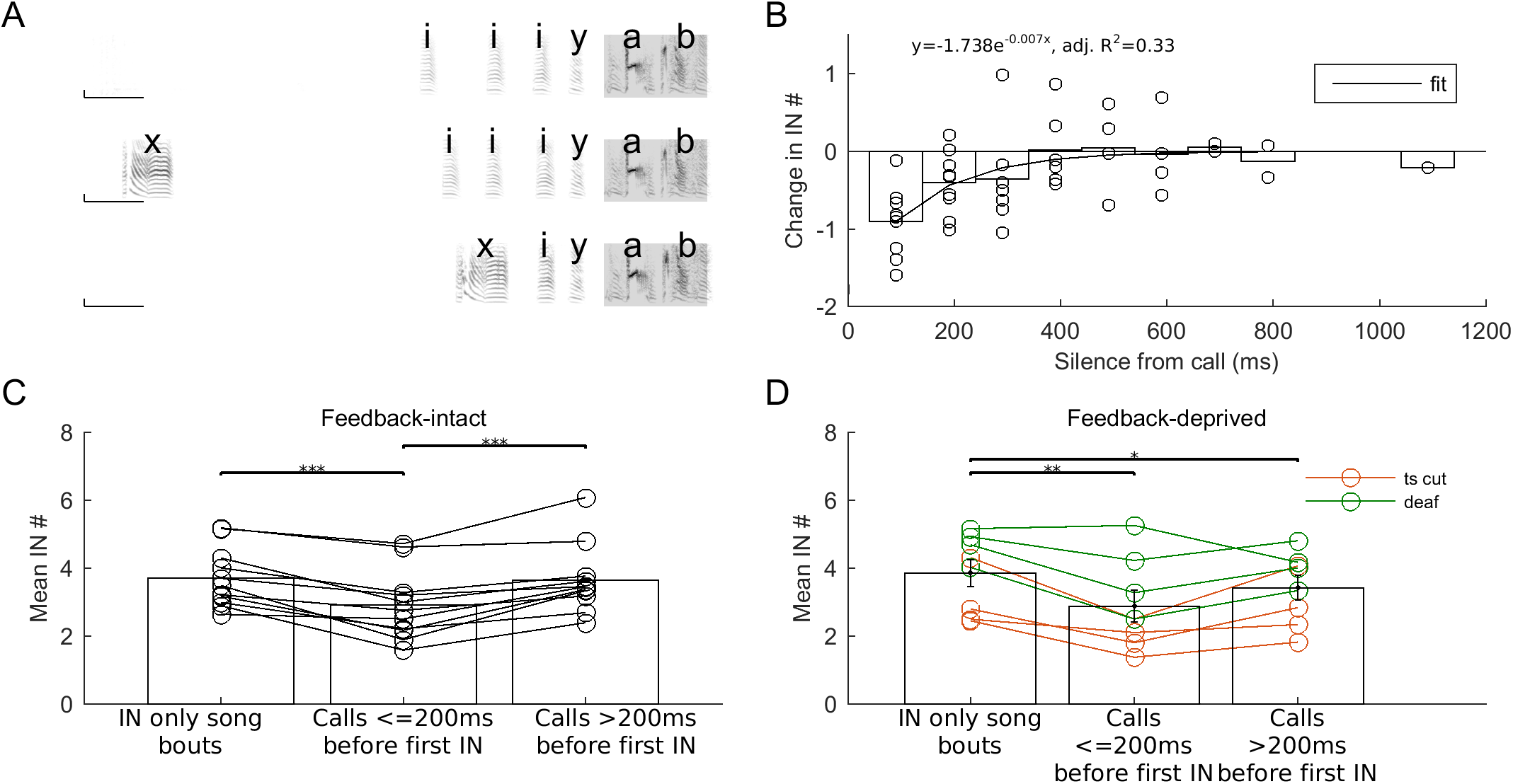
Calls just before the first IN correlate with fewer INs at the start of the bout. (A) Spectrograms of the start of an IN song bout (top) and two call song bouts with the call occuring well before the first IN (middle) or just before the first IN (bottom). ‘i’ and ‘y’ represent INs, ‘a’ and ‘b’ represent motif syllables and ‘x’ represents a call. Scale bar 200ms and 1000 Hz. (B) Silence between the end of the call and the beginning of the first IN vs. change in IN number relative to the mean IN number in IN song bouts. Each circle represents one bird. Bars represent mean across birds and the line represent an exponential fit to the data (y=-1.642e^-0 007x^, adjusted R^2^ = 0.31). (C) and (D) Mean IN number in IN song bouts, call bouts with calls occuring ≤ 200ms before the first IN and call bouts with calls occuring > 200ms before the first IN for unmanipulated, feedback-intact birds (C), and for feedback-deprived birds (D; red – post ts-cut; green – post deafening). Circles represent individual birds and lines connect data from the same bird. Bars and whiskers represent mean and s.e.m. across birds. * represents p < 0.05, ** represents p < 0.01, *** represents p < 0.001, Repeated Measures 1way ANOVA.

**Figure 8.**
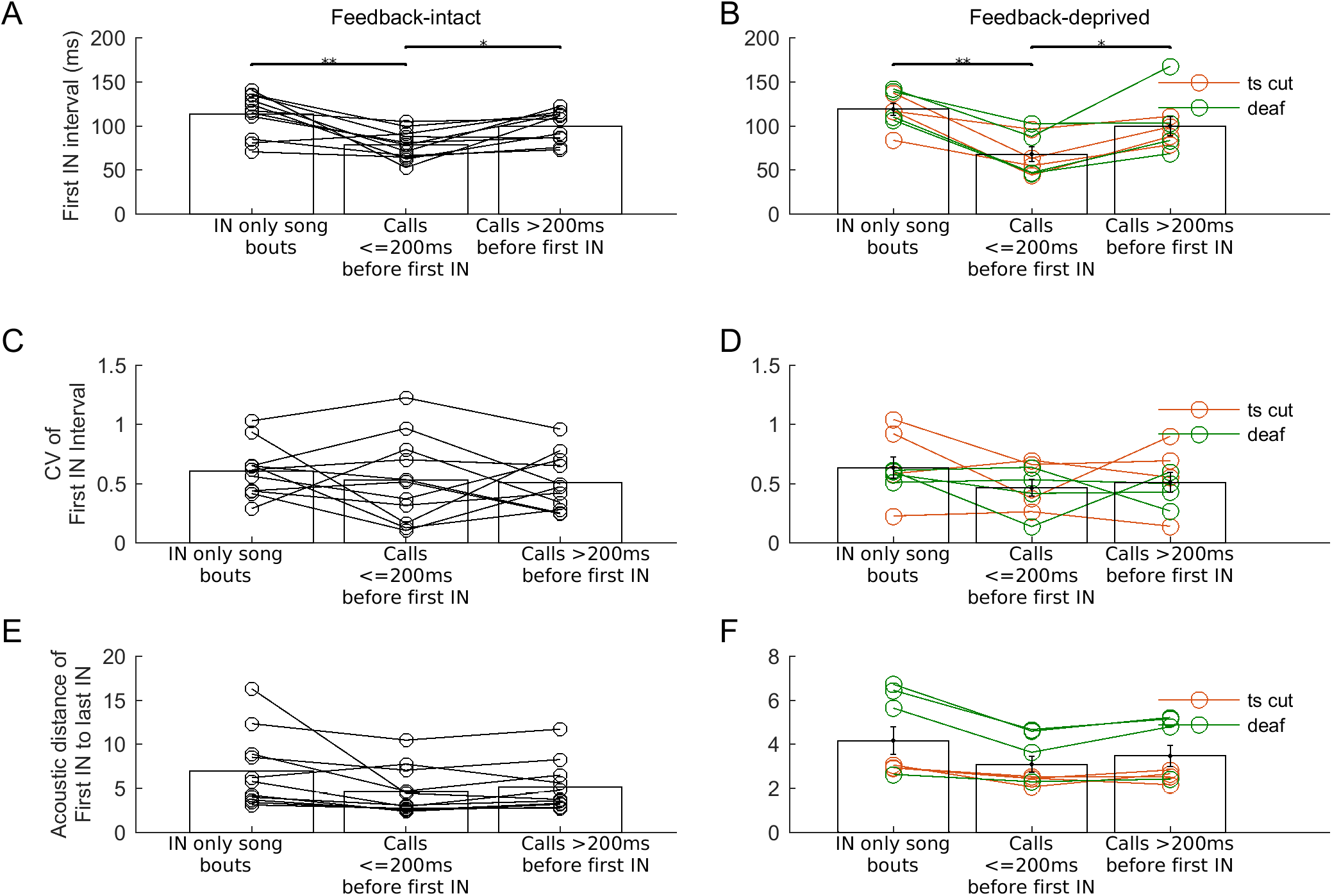
Calls just before the first IN correlate with a shorter interval between first two INs. (A), (B) Mean duration of the interval between the first two INs in IN song bouts, call bouts with calls occuring ≤ 200ms before the first IN and call bouts with calls occuring > 200ms before the first IN for unmanipulated, feedback-intact birds (A), and for feedback-deprived birds (B; red – post ts-cut; green – post deafening). Circles represent individual birds and lines connect data from the same bird. Bars and whiskers represent mean and s.e.m. across birds. (C), (D) Variability of the interval between the first two INs in IN song bouts, call bouts with calls occuring ≤ 200ms before the first IN and call bouts with calls occuring ≤ 200ms before the first IN for unmanipulated, feedback-intact birds (C), and for feedback-deprived birds (D; red – post ts-cut; green – post deafening). Circles represent individual birds and lines connect data from the same bird. Bars and whiskers represent mean and s.e.m. across birds. (E), (F) Mean acoustic distance of the first IN from the last IN in IN song bouts, call bouts with calls occuring < 200ms before the first IN and call bouts with calls occuring > 200ms before the first IN for unmanipulated, feedback-intact birds (E), and for feedback-deprived birds (F; red – post ts-cut; green – post deafening). Circles represent individual birds and lines connect data from the same bird. Bars and whiskers represent mean and s.e.m. across birds. * represents p < 0.05, ** represents p < 0.01, *** represents p < 0.001, Repeated Measures 1way ANOVA.

### Deafening

Deafening was done by bilateral removal of the cochlea under equithesin anesthesia using previously described protocols (Kojima et al., 2013; Konishi, 1965). All of the deaf birds (n=7) were also used in a previous study that examined the effects of deafening on song (Kojima et al., 2013). Here we only analyzed the effects of deafening on introductory note (IN) number and properties. Since, we were interested in the role of real-time auditory feedback on progression from INs to song, we only analyzed IN number and properties for songs recorded 1 day post-deafening.

### Data Analysis

All analysis was done using custom-written scripts in MATLAB.

### Song analysis

Audio files were segmented into syllables based on a user-defined amplitude threshold. Syllables with less than 5ms between them were merged and syllables with duration shorter than 10ms were discarded. Individual syllables were given labels in a semi-automatic manner. They were first assigned labels based on a modified template matching procedure (Glaze and Troyer, 2006) or clustering based on acoustic features calculated using Sound Analysis Pro. Clustering was done using KlustaKwik (https://sourceforge.net/p/klustakwik/wiki/Home/). Labels were then manually checked for all files.

The repetitive sequence or song motif for each bird was identified. Song bouts were defined as groups of vocalizations with atleast one motif syllable that were separated from other such groups by more than 2s of silence (Sossinka and Böhner, 1980). For a subset of birds (n=7 deaf birds; n=6 birds for analysis of call-song bouts and n=7 birds for analysis of day-to-day changes in IN number and properties) with triggered recordings, a number of files did not have 2s of silence before the first vocalization in the file. However, since these were triggered recordings, we assumed that there was silence before the start of the file too and so we considered files with > 0.5s silence at the beginning of file as valid bouts. For a given bird, we used the same criterion before and after surgery to ensure that the criterion did not affect our results. Syllables that were produced in isolation outside of song bouts were identified as calls. All kinds of calls (distance calls, short calls and intermediate calls) (Zann, 1996) were combined together.

As described earlier (Price, 1979; Rajan and Doupe, 2013; Sossinka and Böhner, 1980), syllables that were repeated at the beginning of a bout were considered as introductory notes (INs). Calls were not considered as INs. As described earlier (Zann, 1993), 76.2% of our birds (n=32/42) produced only one IN type. The rest of the birds produced 2 IN types (n=10/42). For all the analysis described, we combined the multiple types of INs together.

For ts-cut birds, syllables and INs lost their characteristic acoustic structure and were reduced to harmonic stacks without any modulation (Fig. 2A middle). However, durations of individual syllables and INs remained same as pre-surgery (Fig. 2A middle, 2B middle, Fig. 5, Supp. Fig. 3). In these birds, syllables were labelled using cluster analysis as described above and INs and motif syllables were matched to pre-surgery INs and motif syllables by examining plots for duration vs. mean frequency for all syllables.

On average, we analyzed 124 song bouts per bird (median = 98 song bouts per bird; range = 11-428 song bouts per bird).

### Temporal and spectral similarity

We quantified changes in song after removal of sensory feedback using temporal and spectral similarity. Temporal similarity was calculated as the maximum of the cross-correlation function between the normalized amplitude envelopes of a pre-surgery template motif and other pre/post surgery motifs (n=9 randomly chosen motifs from pre-surgery and n=10 randomly chosen motifs from post surgery) (Roy and Mooney, 2007). The template motif was proportionally stretched ± 20% to account for differences in duration of the entire motif. As a measure of random temporal similarity between any two zebra finches, we calculated temporal similarity for motifs from 10 random pairs of birds (n=10 motifs each). Spectral similarity (% similarity) was calculated using Sound Analysis Pro (five motifs pre-surgery were compared to each other and to 5 motifs postsurgery) (Tchernichovski et al., 2000). Random spectral similarity was measured for ten random pairs of birds (n=5 motifs each). Song spectral and temporal similarity was calculated only for 5/6 sham surgery birds.

### Characterization of Introductory note (IN) progression

In each bout, the last set of consecutive INs with inter-IN intervals < 500ms, before the first motif syllable were considered for counting IN number in each song bout (Rajan and Doupe, 2013; Sossinka and Böhner, 1980). All of our analysis was restricted to such sequence of INs present at the beginning of each bout.

Intervals between INs were measured as the duration between the end of an IN to the start of the next IN. The first interval was the interval between the first two INs satisfying the above criteria. The last interval was measured as the interval between the last IN and the first motif syllable. As a measure of the progression of IN timing, we quantified the ratio between successive INintervals across all IN sequences. Ratios were averaged across bouts to obtain a mean ratio for each bird. A ratio < 1 indicated a speeding up of successive intervals as shown earlier (Rajan and Doupe, 2013). CV was measured as the standard deviation divided by the mean.

To characterize acoustic properties of INs and their progression to song, we used the acoustic distance to the last IN and the ratio of distance of successive INs respectively. The acoustic distance is an inverse measure of similarity in acoustic properties between an IN and all last INs (Rajan and Doupe, 2013). We calculated 4 acoustic features, namely, duration, log amplitude, entropy and mean frequency for each IN using the matlab code for Sound Analysis Pro (http://soundanalysispro.com/matlab-library). For each day, we randomly chose 50% of the last INs as the reference. The distance of each first IN and the remaining last INs were measured as the Mahalanobis distance of the IN from the reference last INs in the 4-dimensional space formed by the 4 acoustic features. As a measure of acoustic progression of INs, we calculated the ratio of distances of successive INs from the last IN for each IN sequence at the beginning of a bout (50% of the bouts were excluded as the last INs from these bouts were chosen as the reference). A ratio < 1 indicated that successive INs became closer in distance (or more similar) to the last IN, as seen in intact birds (Rajan and Doupe, 2013).

### Analysis of the influence of calls on number and properties of INs

The influence of calls on IN number and properties was analyzed in 16 normal, unmanipulated birds. Bouts where the first IN began < 2000ms after the end of a call were considered as call-song bouts. Bouts with only INs at the beginning were considered as IN song bouts. Birds with a minimum of 5 IN song bouts and 5 call-song bouts were considered for this analysis. For each bird, the number of INs in IN song bouts was subtracted from the number of INs in each call-song bout. For each bird, the change in IN number in call-song bouts was then binned at 100ms resolution starting at 40ms after the end of the call to 1940ms after the end of the call. Across all birds, we fitted an exponential function (matlab fit function) to characterize the dependence of this change in IN number on time between the end of the call and the start of the first IN. Similarly, we also fit exponential functions to the change in the interval between the first two INs and change in acoustic properties of the first IN (Supp. Fig. 4).

For many of the feedback-deprived birds, we did not have enough call-song bouts to carry out a similar analysis. Instead, we divided call-song bouts into two categories: (1) bouts where the first IN started < 200ms after end of the call and (2) bouts where the first IN started > 200ms after the end of the call. 200ms was chosen based on the exponential fit (Fig. 7B) and data availability in the feedback-deprived birds. We calculated IN number, mean and variability of the interval between the first two INs, acoustic distance of the first IN for both these bout categories and compared it with the corresponding properties for IN song bouts (Fig. 7, 8). Only birds with > 3 call song bouts in both of these categories were considered for analysis. Further, we combined ts-cut and deaf birds as our previous results showed that both manipulations had no effect on IN number and properties.

### Statistics

Wilcoxon signed rank test was used for paired comparisons of temporal similarity (Fig. 2C), spectral similarity (Fig. 2D), changes in IN/motif syllable acoustic features (Fig. 3 and Supp. Fig. 1), mean IN number (Fig. 4D), IN number CV (Fig. 4E) and progression in IN features (Supp. Fig. 3). For comparing progression in IN timing and IN acoustic structure after removal of feedback (Fig. 5D, 5E and Fig. 6D), we used Repeated Measures 2-way ANOVA using IN position (First vs. Last) as one factor and Time (Pre surgery vs. Post surgery) as the second factor (Matlab code from https://in.mathworks.com/matlabcentral/fileexchange/6874-two-way-repeated-measures-anova). For comparing IN number and properties in bouts where calls preceded the first IN, we used repeated measures 1-way ANOVA (Fig. 7C, 7D and Fig. 8). If the ANOVA p-value was < 0.05, we used a post-hoc Tukey Kramer test to identify groups that were significantly different (Fig. 7C, 7D and Fig. 8). For comparing changes in IN number and properties after surgery with day-to-day changes, we used Kruskal-Wallis ANOVA (Supp. Fig. 2). Pearson’s correlation coefficient was used to assess the correlation between first IN properties and time-to-song initiation (Fig. 9).

**Figure 9.**
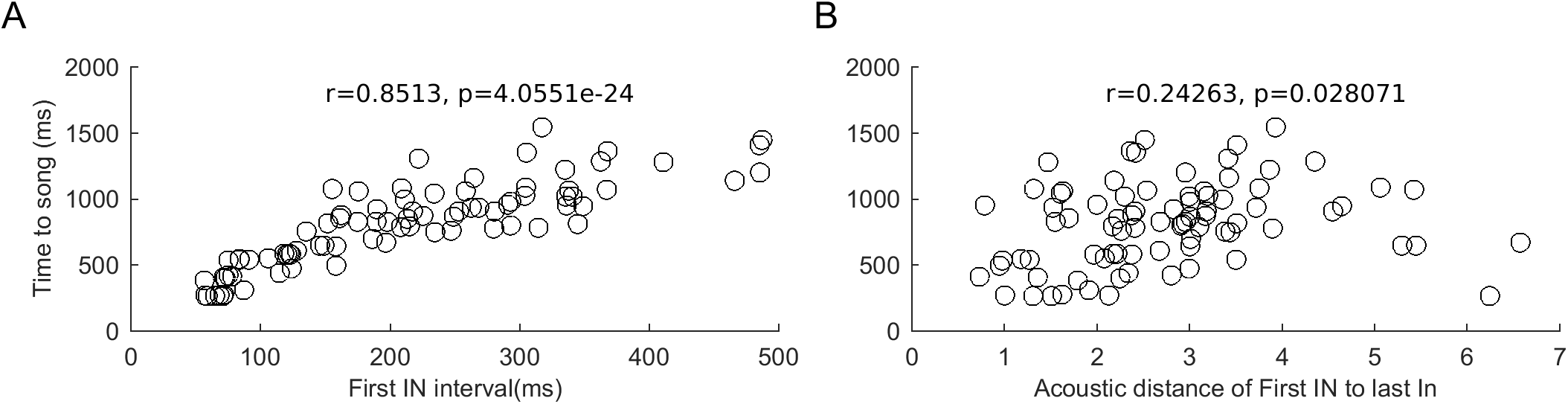
“Initial” state of IN progresison correlates with time to song initiation. (A) and (B) Correlation between the interval between the first two INs (A) or the acoustic distance of the first IN to the last IN (B) and the time to start of song (onset of the first motif syllable) for one bird. Circles represent data from individual bouts.

All of the tests used and the associated p-values are provided in Supp. Table 1. A significance level of p = 0.05 was used throughout.

## RESULTS

To see if INs provide sensory feedback for song initiation (Fig. 1B), we analyzed the number and progression of INs after removal of either proprioceptive (n=8 birds) or auditory feedback (n=7 birds). As a control, sham surgeries were performed in a separate group of birds (n=6). Since, we were interested in self-initiated movement sequences, we focused on undirected songs produced when the bird was alone.

### Song temporal structure was not affected after removal of auditory or proprioceptive feedback but song spectral structure was disrupted after removal of proprioceptive feedback

Proprioceptive feedback was removed by bilaterally cutting the tracheosyringeal nerve (Bottjer and Arnold, 1984; Vicario, 1991; Williams and McKibben, 1992) (ts-cut; n=8 birds, see Methods) and auditory feedback was eliminated by bilateral removal of the cochlea (Konishi, 1965) (deaf; n=7 birds, see Methods). While our main focus was on INs, we first compared both the spectral and temporal structure of songs produced soon after surgery (1-10 days post surgery – see Methods) with songs produced just before surgery. Songs of birds subjected to sham surgery remained similar to pre-surgery songs in both spectral and temporal structure (Fig. 2A, 2B, 2C, 2D). Birds subjected to either deafening or ts-cut showed changes in both spectral structure and temporal structure (Fig. 2A, 2B, 2C, 2D; p < 0.05, Wilcoxon sign-rank test). Similar to earlier studies (Konishi, 1965; Price, 1979), these changes were small in deaf birds. Both the spectral (Fig. 2D) and temporal structure (Fig. 2C) of songs produced soon after deafening remained more similar than expected by chance to pre-deafening songs. In ts-cut birds, song spectral structure was completely disrupted after nerve cut and songs were no longer similar to pre-surgery songs (Fig. 2D). The tracheosyringeal nerve contains both efferent and afferent nerves carrying motor input to the syringeal muscles and proprioceptive feedback from the syringeal muscles respectively (Bottjer and Arnold, 1984). Since the nerve cut also affected efferent nerves carrying neural input to the syringeal muscles, spectral structure of song was disrupted as reported earlier (Roy and Mooney, 2007; Vicario, 1991; Williams and McKibben, 1992). Specifically, all syllables lost their characteristic acoustic structure and were reduced to harmonic stacks (Fig. 2A middle). However, the temporal structure remained more similar, than expected by chance, to songs produced before nerve cut (Fig. 2B middle and Fig. 2C) since motor input to the respiratory muscles was not affected in ts-cut birds (Bottjer and Arnold, 1984; Roy and Mooney, 2007; Vallentin and Long, 2015; Vicario, 1991; Williams and McKibben, 1992). Thus, consistent with earlier studies (Bottjer and Arnold, 1984; Roy and Mooney, 2007; Vallentin and Long, 2015; Vicario, 1991; Williams and McKibben, 1992), we also found that ts-cut altered both proprioceptive and auditory feedback, while deafening disrupted only auditory feedback.

### IN acoustic structure, not duration, is affected by removal of proprioceptive feedback

We next quantified changes to the acoustic structure of INs after removal of either proprioceptive or auditory feedback. Similar to changes in song syllable structure (Fig. 2, Supp. Fig. 1, middle column), INs also became harmonic stacks after surgery in ts-cut birds as seen by increased pitch goodness and decreased frequency modulation (Fig. 2A middle, Fig. 3, p < 0.05, Wilcoxon sign-rank test). Despite this change, we could identify INs because IN duration, mean frequency and amplitude did not change significantly after surgery (Fig. 3A middle – see Methods for details of IN identification procedure in ts-cut birds). The position of INs at the beginning of the bout was also maintained (Fig. 2A middle). Post-deafening, INs were softer and had reduced mean frequency (Fig. 3B, 3D right, p < 0.05, Wilcoxon sign-rank test). However, song syllables were also softer after deafening (Supp. Fig. 1, right column, p < 0.05, Wilcoxon sign-rank test) suggesting that these changes could have been a result of a change in microphone position after surgery. No significant changes in IN acoustic structure were seen after sham surgery (Fig. 3, left, p > 0.05, Wilcoxon sign-rank test).

### Mean IN number before song was not affected by removal of proprioceptive or auditory feedback

We next analyzed the mean and variability of IN number before each song (see Methods). Mean number of INs before song (Fig. 4A, 4B, 4C, 4D) and the variability in IN number (measured by the CV – Fig. 4E) did not change significantly soon after surgery in sham-surgery, ts-cut and deaf birds (Fig. 4D and 4C, p > 0.05, Wilcoxon sign-rank test). Infact, changes in mean IN number post-surgery for feedback-deprived birds were not different from day-to-day changes in IN number seen in normal, unmanipulated, birds (Supp. Fig. 2A). This further strengthened our conclusion that IN number was unaffected by removal of proprioceptive or auditory feedback.

### Progression of IN timing to song is not affected by removal of proprioceptive of auditory feedback

We have previously shown that progression of INs to song is accompanied by changes in both timing and acoustic structure of INs within a bout (Rajan and Doupe, 2013). We first considered IN timing. Specifically, intervals between successive INs progress from a longer, more variable, first interval to a shorter, more stereotyped, interval between the last IN and song (Rajan and Doupe, 2013). This progression in IN timing was unchanged after surgery in sham surgery, ts-cut and deaf birds (Fig. 5A, 5B, 5C). After surgery, the interval between the first two INs remained longer and more variable than the interval between the last IN and song in ts-cut, deaf and sham surgery birds (Fig. 5D, 5E; p < 0.05 for First vs. Last, Repeated Measures 2-way ANOVA). Importantly, removal of feedback did not alter either the mean or variability of both the first interval and the last interval (Fig. 5D, 5E; p > 0.05 for Pre vs. Post, Repeated Measures 2-way ANOVA). A number of other aspects of IN timing were also not affected by removal of auditory or proprioceptive feedback and changes in IN timing post-surgery were similar to day-to-day changes seen in unmanipulated birds (Supp. Fig. 2B, 2C, 2D, Supp. Fig. 3A – see Methods). These results showed that the timing of INs and their progression did not depend on intact sensory feedback.

### Progression of IN acoustic features to song is not affected by removal of proprioceptive of auditory feedback

Similar to IN timing, the acoustic structure of INs has also been shown to progress to a consistent last-IN state just before song (Rajan and Doupe, 2013). Although individual INs in each bout look very similar (Fig. 6A, 6B, 6C – top), we have previously shown that the first IN is less similar to the last IN across bouts. We quantified this by calculating the similarity between the first IN and the last IN before and after surgery (acoustic distance to last IN – smaller the distance, greater the similarity and vice versa; see Methods; see Fig. 6A, 6B, 6C for representative examples for sham surgery, ts-cut and deaf birds). Since we were interested in the progression, we calculated similarity to the last IN on the same day (pre-surgery last IN for presurgery and post-surgery last IN for post-surgery; see Methods). For each day, half of the last INs across all bouts were randomly chosen as a reference. The rest of the last INs and all of the first INs were then compared to this reference using the acoustic distance as an inverse measure of similarity (see Methods). The first IN was significantly different from the last IN (larger distance – Fig. 6D, p = 0.0544 for First vs. Last in ts-cut and p < 0.05 for First vs. Last in deaf birds, Repeated Measures 2-way ANOVA) before and after surgery in ts-cut and deaf birds. However, this difference was smaller in sham-surgery birds both before and after surgery and did not reach significance (Fig. 6D left, p = 0.2469 for First vs. Last in sham-surgery birds, Repeated Measures 2-way ANOVA). Importantly, in all groups of birds, removal of feedback did not affect any of the measures of progression (p > 0.05, Pre vs. Post, Repeated Measures 2-way ANOVA). These results showed that INs still progressed from a first IN that was significantly different from the last IN to a more consistent last IN even in the absence of auditory or proprioceptive feedback. A number of other aspects of IN acoustic structure progression to song were also not affected by removal of auditory or proprioceptive feedback and remained similar to day-to-day changes seen in unmanipulated birds (Supp. Fig. 2E, 2F, 2G and Supp. Fig. 3B – see Methods). As mentioned earlier, ts-cut birds lacked neural input to the syringeal muscles in addition to the loss of proprioceptive feedback from the syringeal muscles. The continued progression of IN acoustic features suggested that this progression is a result of changing respiratory drive, since neural input to the respiratory muscles remained intact in these birds.

Overall, these results show that IN number and progression of IN timing are not dependent on intact sensory feedback (auditory and proprioceptive) and suggest that INs reflect an internal neural preparatory process (Fig. 1C).

### IN number was reduced when calls preceded the first IN of a song bout

We have previously shown changes in neural activity in premotor nucleus HVC, starting hundreds of milliseconds before the start of the first IN of an undirected song bout (Rajan, 2018). While the origin of these changes is not known, these changes are thought to represent internal preparatory activity. Our current results suggested that INs also reflected internal neural preparatory processes, possibly a continuation of preparatory activity that began well before the first IN. If this was the case, we predicted a correlation between the number of INs and the levels of preparatory activity that preceded it. In our current study, we did not record neural activity to directly test this prediction. However, we behaviorally tested this prediction, in a separate set of unmanipulated birds (n=16), by examining the number of INs in song bouts where calls (other non-song vocalizations) preceded the first IN (call-song bouts – see Methods).

Calls are partially learned or unlearned vocalizations that are acoustically distinct from song and are initiated by separate neural pathways (Simpson and Vicario, 1990; Vicario, 2004; Zann, 1996). Many aspects of calls are controlled by song motor nuclei and increased neural activity is seen in many of the song motor nuclei before and during calls (Benichov et al., 2016; Danish et al., 2017; Hahnloser et al., 2002; Kozhevnikov and Fee, 2007; Long and Fee, 2008; Rajan, 2018; Simpson and Vicario, 1990; Vyssotski et al., 2016; Yu and Margoliash, 1996). Further, we have previously shown the presence of higher levels of preparatory activity in HVC before the first IN when calls precede the first IN of a song bout (Rajan, 2018). Therefore, we expected fewer INs when the first IN of a song bout was preceded by calls.

Calls occured at variable times before the first IN in a small fraction of bouts (Fig. 7A; mean +/− s.e.m. of interval between end of call and start of first IN = 471.2 +/− 48 ms; mean +/− s.e.m. of CV of interval between end of call and start of first IN = 0.86 +/− 0.07; n=16 birds). Consistent with our prediction of fewer INs in call song bouts, we observed fewer INs when calls occurred before the first IN (n=16 birds; mean +/− s.e.m. for IN song bouts = 3.7 +/− 0.24, mean +/− s.e.m. for call song bouts = 3.4 +/− 0.25, p = 0.03, Wilcoxon sign-rank test). This reduction was dependent on the time between the end of the call and the start of the IN; shorter the time, greater the reduction (Fig. 7B, p < 0.05, adjusted R^2^ = 0.32 for an exponential fit, see Methods). In both feedback-intact and feedback-deprived birds, song bouts where the first IN began < 200ms after the end of a call, had fewer INs when compared to song bouts with only INs at the beginning or song bouts where the first IN began > 200ms after the end of the call (Fig. 7C, 7D, p < 0.05, repeated measures 1-way ANOVA and post-hoc Tukey Kramer test). These results showed that the presence of calls just before the first IN of a song bout correlated with fewer INs in both feedback-intact and feedback-deprived birds and suggest that INs could represent a continuation of on-going neural preparatory activity that begins well before the first IN (Rajan, 2018).

### Calls just before the first IN of a song bout correlated with altered “initial” state

Given that both IN timing and acoustic features progress towards a consistent “ready” state just before song, we hypothesized that calls might reduce IN number by speeding up this progression. Consistent with this idea, song bouts where the first IN began < 200ms after the end of call had a significantly shorter interval between the first 2 INs when compared to song bouts with only INs or song bouts where the first IN began > 200ms after the end of the call (Fig. 8A, 8B; p < 0.05, Repeated Measures 1-way ANOVA followed by post-hoc Tukey-Kramer test). This was true both in feedback-intact (Fig. 8A) and feedback-deprived birds (Fig. 8B). In feedback-intact birds, the decrease in interval between the first two INs in bouts with calls was correlated with the time between the end of the call and the start of the first IN, though the strength of the correlation was moderate (Supp. Fig. 4A, Adjusted R^2^ for exponential fit = 0.15). In contrast to the changes in IN timing, neither the variability of the interval between the first two INs nor the acoustic structure of the first IN showed any differences based on whether calls were present before the first IN or not (feedback-intact birds - Fig. 8C, 8E; feedback-deprived birds – 8D, 8F; p > 0.05, Repeated Measures 1-way ANOVA). However, in feedback-intact birds, relative to bouts with only INs, the acoustic structure of the first IN after a call was more similar to the last IN (Supp. Fig. 4B). The change in acoustic structure was correlated with the time between the end of the call and the start of the first IN, but the strength of the correlation was weak (Supp. Fig. 4B, Adjusted R^2^ for exponential fit = 0.09). Overall, these results showed that the presence of calls correlated with a change in IN timing (shorter interval between the first two INs) potentially causing the reduction in IN number before song.

### “Initial” state of IN progression correlated with time to song initiation

Our results suggested that INs and the progression of IN timing and acoustic features represent on-going neural preparation. In other systems, neural preparatory activity is strongly correlated with the time to movement initiation – greater the progress of preparation, shorter the time to movement initiation (Churchland et al., 2006a; Shenoy et al., 2011; Shenoy et al., 2013). Similar to this, we found a significant correlation between the length of the interval between the first two INs and the time to song initiation in all birds (Fig. 9A, one bird; n=16/16 birds; mean r = 0.77; range = 0.57 – 0.90). How similar the first IN was to the last IN was also correlated with the time to song initiation, albeit to a weaker extent in 14/16 birds (Fig. 9B, one bird; n=14/16 birds; mean r = 0.32; range = −0.39 – 0.62). These data strengthened our conclusions that INs and their progression reflect on-going internal neural preparatory processes and suggested that IN timing and acoustic features reflect the “current” state of neural preparation.

## DISCUSSION

In this study, we show that real-time auditory and/or proprioceptive feedback is not required for initiation of adult zebra finch song. We also show that the progression of INs, the repeated pre-song vocalizations, from a variable initial state to a more stereotyped final state is also independent of real-time sensory (auditory and/or proprioceptive) feedback. Further, we show, in both feedback-intact and feedback-deprived birds, that fewer INs are present when the first IN of a song bout occurs within 200ms of the end of a call (other non-song vocalization). In such cases, IN timing was closer to the final state. Finally, the “initial” state of IN progression was correlated with the time to song initiation. Overall, these results demonstrate that INs do not provide real-time sensory feedback for the motor system to prepare for song initation. Rather, INs might simply reflect a continuation of internal neural preparatory process with timing and acoustic properties of INs reflecting the “current” state of this preparation.

### Contributions of respiratory feedback to song initiation

One feedback that we did not alter is respiratory feedback from the air sacs (Méndez et al., 2010). However, previous work strongly suggests that respiratory feedback may not contribute to IN initiation. First, one earlier study showed that disrupting respiratory pressure during short syllables (of the order of 60ms) did not disrupt song progression (Amador et al., 2013) Given that INs are short syllables of the order of 60ms, INs may not require real-time respiratory feedback for progression to the next syllable (or song). Second, unilateral disruptions of vagal feedback mostly affected syllables at the end of song (Méndez et al., 2010). Finally, sparse, patterned neural activity of one class of neurons in premotor nucleus HVC during singing was also not affected by removal of sensory feedback including respiratory feedback (Vallentin and Long, 2015). All of these data suggest that respiratory feedback does not play a role in IN progression.

### Long-term requirement for sensory feedback

Song production in adult birds does not depend on real-time sensory feedback (Bottjer and Arnold, 1984; Konishi, 1965) and our results show that song initiation also does not depend on real-time sensory feedback. However, long-term song maintenance does require intact sensory feedback as shown by song degradation starting many weeks after deafening (Horita et al., 2008; Nordeen and Nordeen, 1992; Williams and McKibben, 1992). Similarly, it is possible that sensory feedback could be necessary in the longer term for maintenance of IN progression to song (our study focused on songs produced within 10 days of removal of feedback). It would also be interesting to see if song degradation seen at later time-points after deafening is linked to (or caused by) a change in IN progression to song. If INs represent preparatory vocalizations, such a link would be expected as small changes in the neural preparatory state in primates is correlated with changes in features of the upcoming movement (Afshar et al., 2011; Churchland et al., 2006b).

### Comparison of INs to motor preparation in other systems

Preparatory neural activity has been described as a slow change in neural activity, starting as early as 1 second before the start of a movement (Chen et al., 2017; Churchland et al., 2006b; Gao et al., 2018; Lee and Assad, 2003; Li et al., 2015; Maimon and Assad, 2006; Murakami et al., 2014; Romo and Schultz, 1987; Tanji and Evarts, 1976). One important characteristic of this preparatory activity appears to be a decrease in variability across trials (Churchland et al., 2006a; Churchland et al., 2006c). The decrease in variability as INs progress to song (Rajan and Doupe, 2013) is very similar to the decrease in variability in neural activity seen before the start of a movement. Together with our current data showing that sensory feedback is not important for progression of INs to song, these results suggest that INs may represent preparatory activity. Additionally, earlier studies have shown the presence of preparatory neural activity in song control areas well before the first IN of undirected song bouts (Hessler and Doupe, 1999; Kao et al., 2008; Rajan, 2018; Woolley et al., 2014) and directed song bouts (Daliparthi et al., 2018 preprint). Thus, INs may reflect a continuation of this preparatory activity that begins hundreds of milliseconds before the first IN.

### Overt movements in other systems as motor preparation

Our results suggest that overt vocalizations (INs) serve as preparatory activity. Previous studies describing neural preparatory activity, in primates and rodents, before the onset of a movement have not described similar overt movements as motor preparation (Chen et al., 2017; Churchland et al., 2006a; Gao et al., 2018; Murakami et al., 2014; Romo and Schultz, 1987; Tanji and Evarts, 1976). However, all of these studies have involved training animals to perform a task and animals are rewarded for maintaining stable posture without movements until a “GO” signal is provided for movement initiation. Therefore, overt preparatory movements, if present during the early stages of learning, would not be reinforced. This raises two interesting predictions for further experiments. (1) Are overt movements present at early stages of learning in primates and rodents too? (2) Songbirds learn their song with internal reinforcement cues that only reinforce similarity to the tutor song (or tutor song memory) (Fee and Scharff, 2010). It would be interesting to see if INs are learned similar to song learning. Additionally, there are human studies showing the presence of small eye movements (microsaccades) and small limb movements while waiting for a “GO” cue to perform an eye or limb movement (Betta and Turatto, 2006; Cohen and Rosenbaum, 2007; Corneil and Munoz, 2014). Changes in pupil size have also been shown to correlate with preparatory activity (Wang et al., 2015). This suggests that overt movements like INs may be more common before the start of naturally learned movements and may reflect motor preparation.

### Mechanisms for IN progression to song

Our results show that sensory feedback is not essential for IN progression to song. Rather, the properties of INs correlate with “time to song initiation” and may reflect the “current” state of motor preparation. How do the properties of INs change to progress to song? In our current study, we showed that the presence of calls prior to the first IN was correlated with shorter intervals between the first two INs and fewer INs before song. Similarly, shorter intervals between the first two INs has also been observed when neural preparatory activity in premotor nucleus HVC, precedes the first IN (Rajan, 2018). Since calls are also associated with increased neural activity in many song control areas (Benichov et al., 2016; Danish et al., 2017; Hahnloser et al., 2002; Kozhevnikov and Fee, 2007; Vyssotski et al., 2016; Yu and Margoliash, 1996), the intervals between successive INs may reflect a history of increased activity within these interconnected motor regions. The shorter interval might also lead to short-term plasticity that might facilitate song initiation by speeding up IN progression. Such short-term plasticity has been observed in the inputs to premotor nucleus HVC (Coleman et al., 2007). Further experiments disrupting short-term plasticity or disrupting activity in motor control regions during IN production could help understand the mechanisms of IN progression to song.

Independent of the mechanisms, our results suggest that simple movements, like INs, that precede the initiation of learned motor sequences, like song, reflect ongoing internal neural preparatory processes.

## Acknowledgements

We would like to thank Prakash Raut for help with bird colony maintenance. We would like to thank Michael Long, Hamish Mehaffey, Anand Krishnan, Deepa Subramanyam, Girish Deshpande and members of the Rajan and Krishnan Labs for useful discussions and comments on the manuscript.

## Competing Interests

The authors declare no competing or financial interests.

## Author contributions

DR and RR designed the experiments. DR performed the experiments. SK performed the deafening experiments. DR and RR analyzed the data. DR and RR wrote the paper with inputs from SK.

## Funding

This work was supported by a Department of Biotechnology (DBT) Ramalingaswami Fellowship (BT/HRD/35/02/2006) and a grant from the Department of Science and Technology (EMR/2015/000829) to RR. We would also like to acknowledge travel support from DBT, CTEP and the Infosys Foundation to DR. The deafening experiments, carried out at UCSF, San Francisco, were supported by an NIH R01 (MH55987) to Allison Doupe.

## SUPPLEMENTAL INFORMATION

Supplemental information includes 4 supplemental figures and 1 supplemental table.

## SUPPLEMENTARY INFORMATION SUPPLEMENTARY FIGURE LEGENDS

**Supplementary Figure 1.**
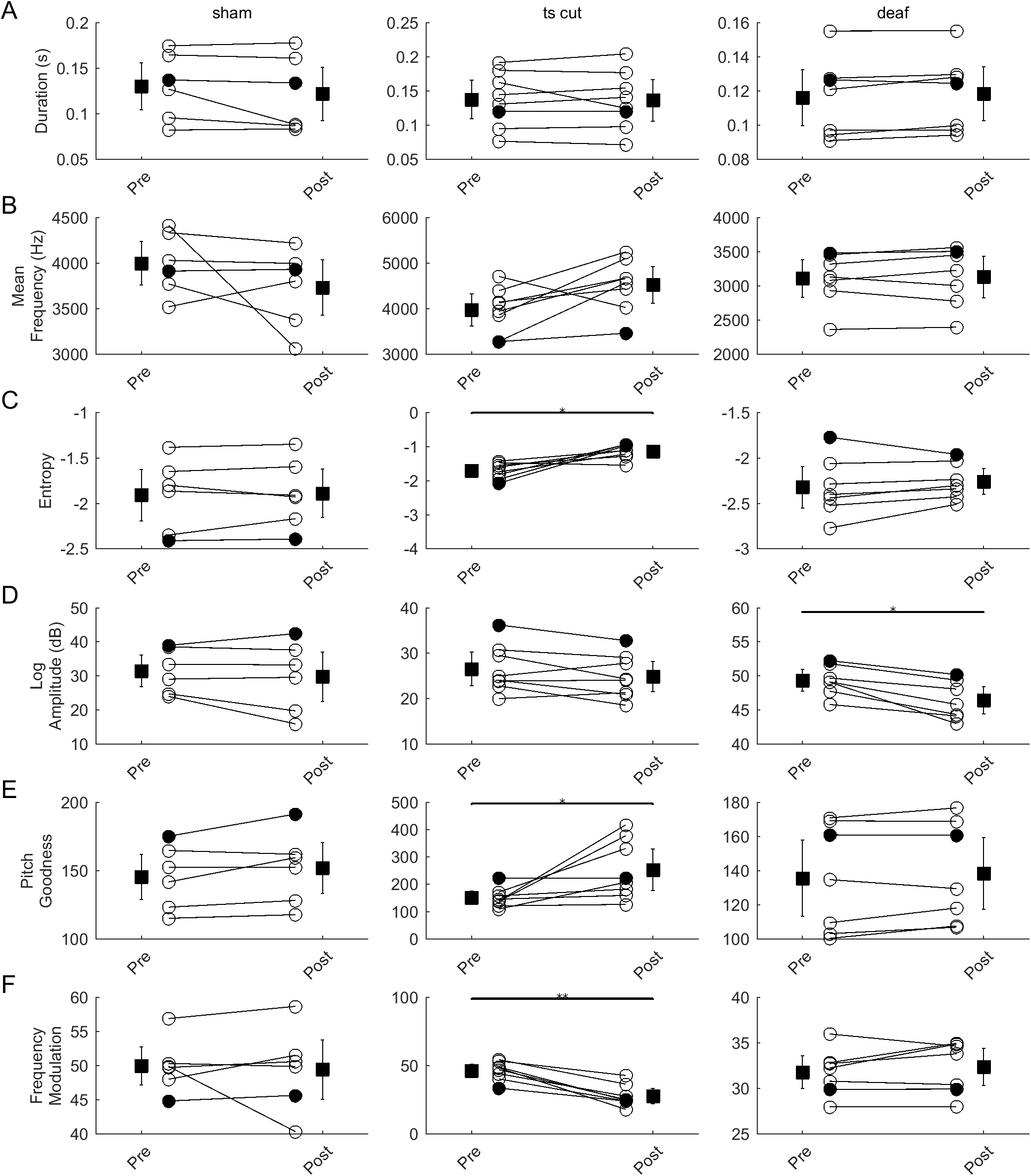
Changes in motif syllable acoustic features after removal of sensory feedback. (A), (B), (C), (D), (E) and (F) Acoustic properties of motif syllables pre and post sham surgery (left column), ts-cut surgery (middle column) or deafening (right column). Circles represent average across all motif syllables for individual birds and lines connect data from the same bird. Squares and whiskers represent mean and s.e.m. across birds. Acoustic features plotted are duration (A), mean frequency (B), entropy (C), log amplitude (D), pitch goodness (E) and frequency modulation (F). * represents p < 0.05, ** represents p < 0.01, Wilcoxon sign-rank test.

**Supplementary Figure 2.**
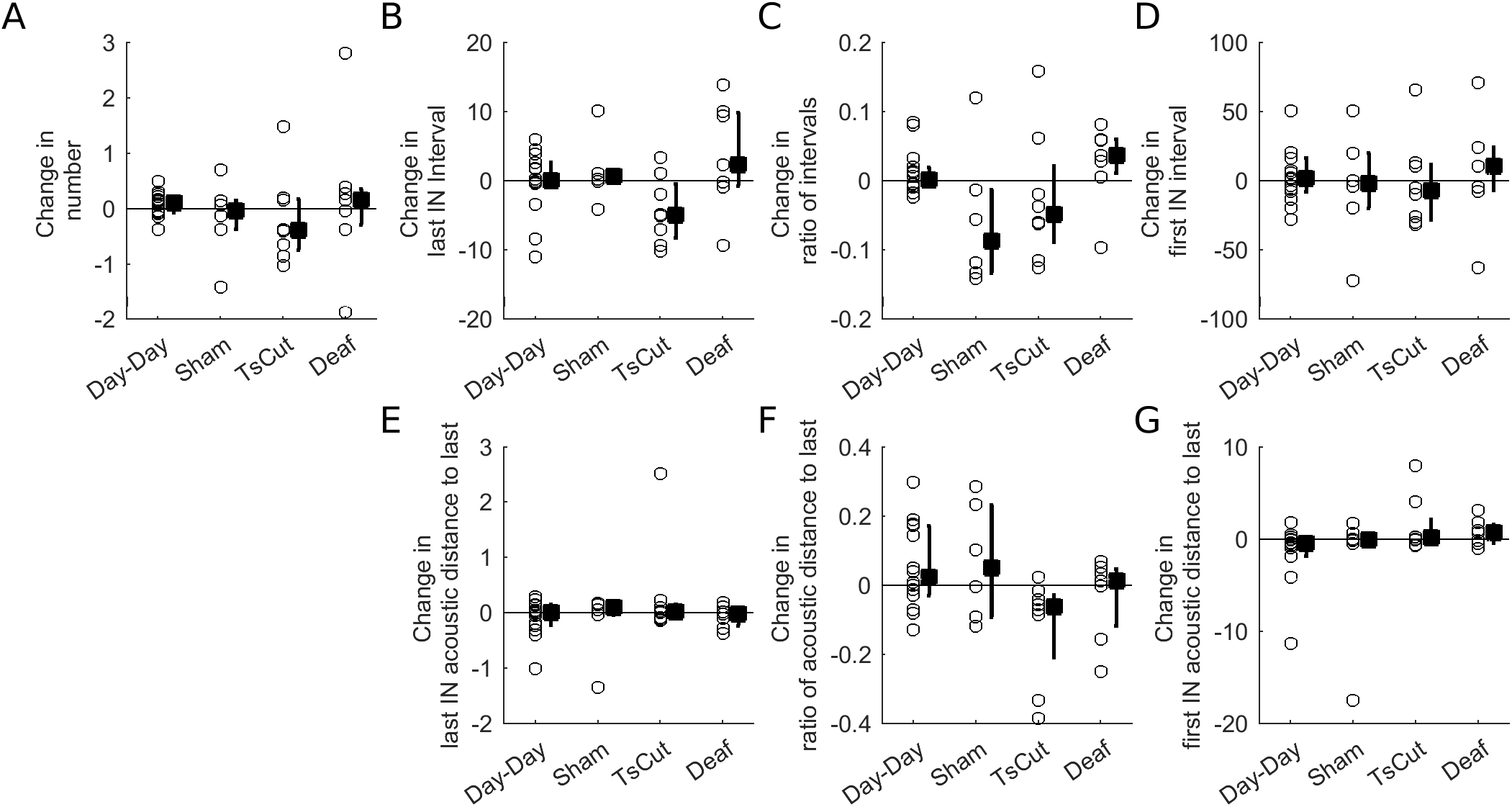
Comparison of day-to-day changes in IN properties with changes observed after surgery for sham-surgery, ts-cut and deaf birds. (A), (B), (C), (D), (E), (F) and (G) Comparison of change in IN properties before and after sham-surgery, ts-cut surgery or deafening with day-to-day changes in IN properties. Changes in IN number (A), Last IN interval (B), Ratio of successive intervals (C), first IN interval (D), last IN acoustic distance to last IN (E), ratio of acoustic distances of successive INs to the last IN (F) and first IN acoustic distance to the last IN (G) are plotted. Circles represent individual birds. Squares and whiskers represent median and inter-quartile range across all birds.

**Supplementary Figure 3.**
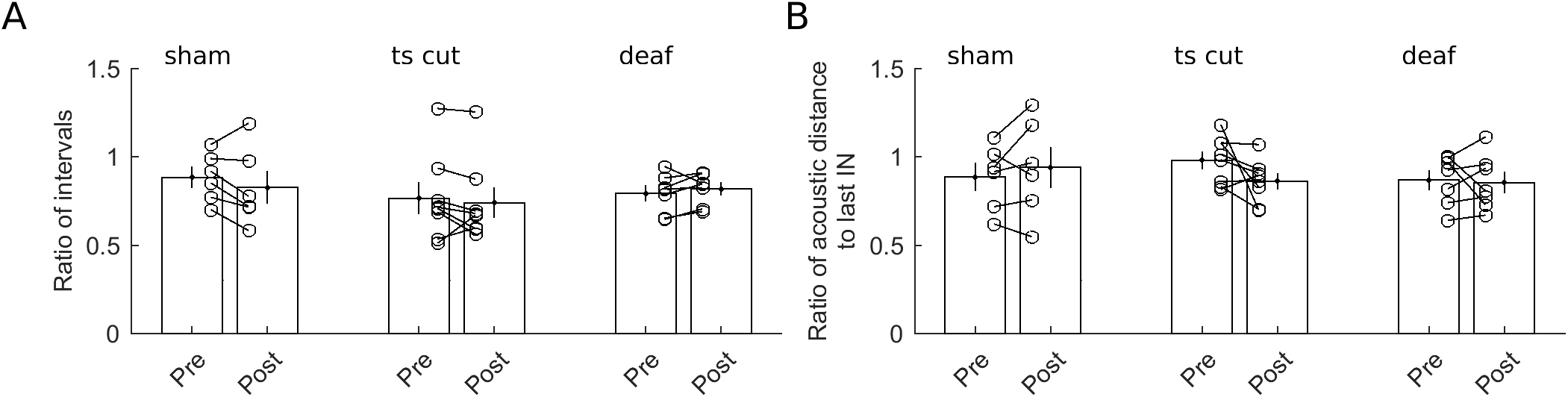
Progression of IN timing and acoustic features is not altered by removal of auditory or proprioceptive feedback. (A), (B) Ratio of successive inter-IN intervals (A) and ratio of acoustic distances of successive INs from the last IN (B) for all birds before and after sham-surgery, ts-cut surgery or deafening. Each circle represents data from an individual bird and lines connect data from the same bird. Bars and whiskers represent mean and s.e.m. across birds

**Supplementary Figure 4.**
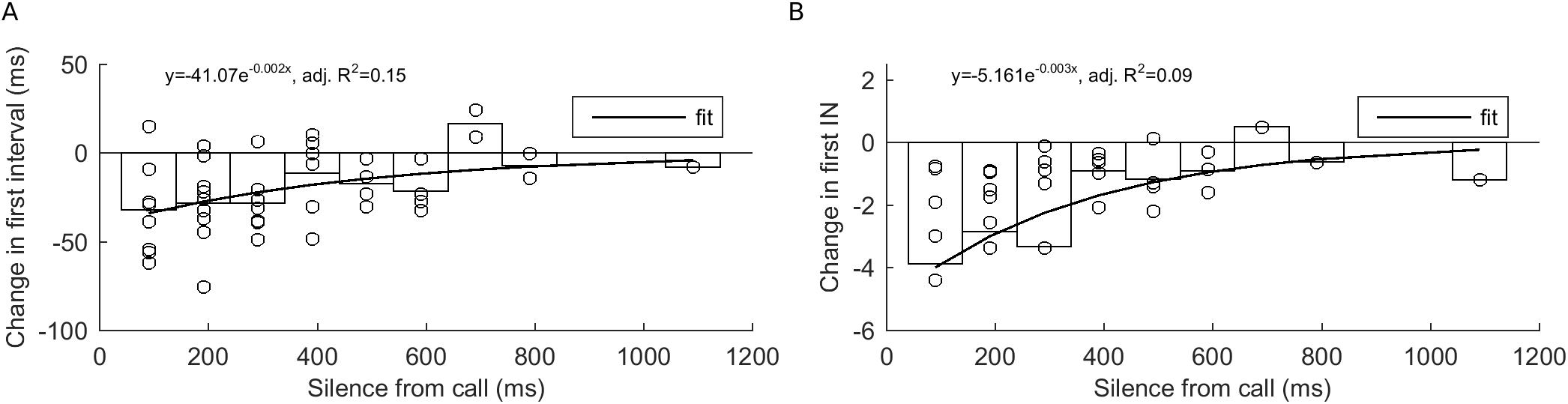
Changes in IN “initial” state in bouts with calls before the first IN also depend on the time between the end of the call and the start of the first IN. (A) Silence between the end of the call and the beginning of the first IN vs. change in interval between the first two INs in call song bouts relative to the interval between the first two INs in IN song bouts. Each circle represents one bird. Bars represent mean across birds and the line represent an exponential fit to the data (y=-41.969e^-0.002x^, adjusted R^2^ = 0.16). (B) Silence between the end of the call and the beginning of the first IN vs. change in acoustic distance of the first IN to the last IN in call song bouts relative to acoustic distance of the first IN from the last IN in IN song bouts. Each circle represents one bird. Bars represent mean across birds and the line represent an exponential fit to the data (y=-1.873e^-0 002x^, adjusted R^2^ = 0.09).

**Supplementary Table 1.**
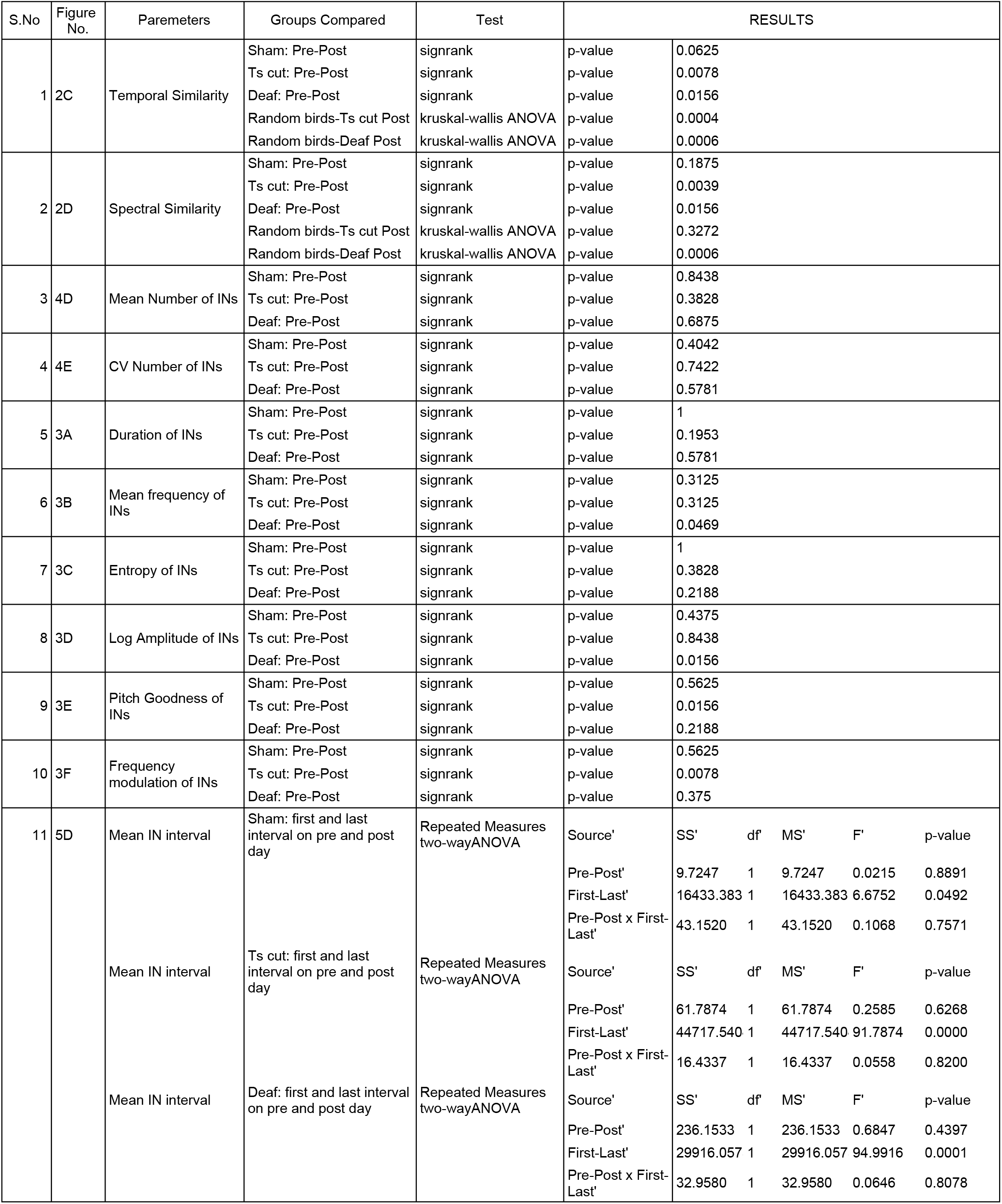
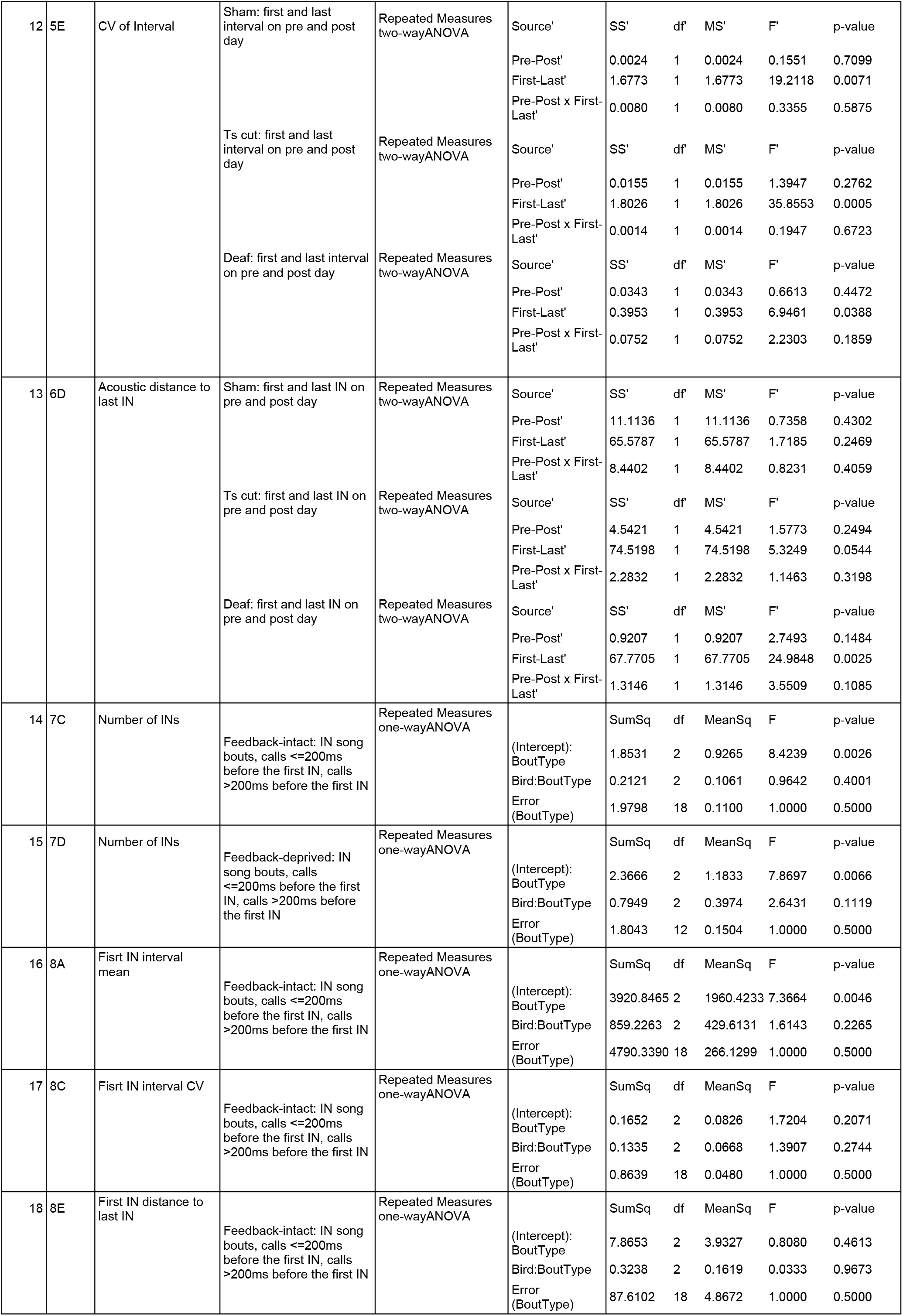
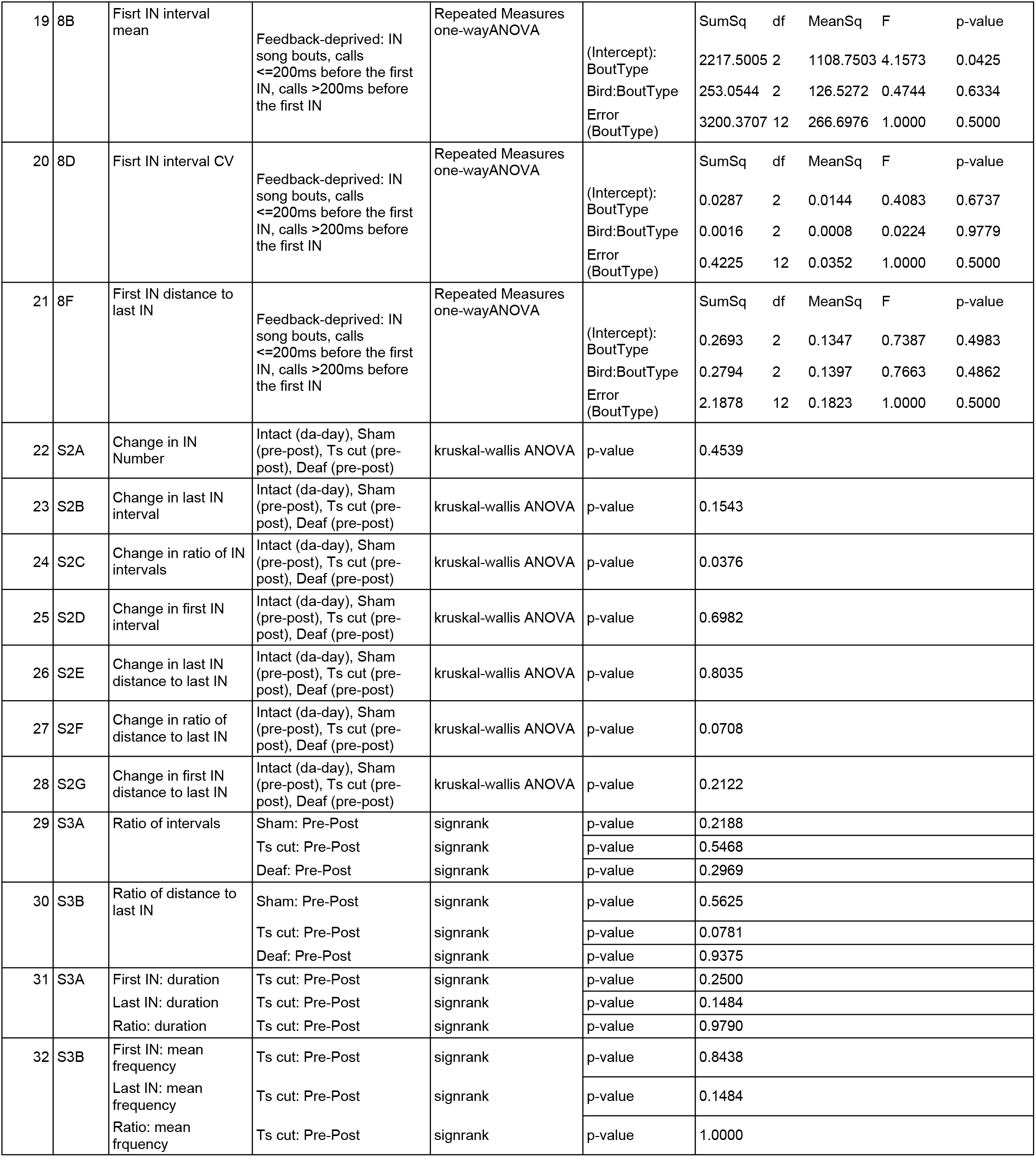
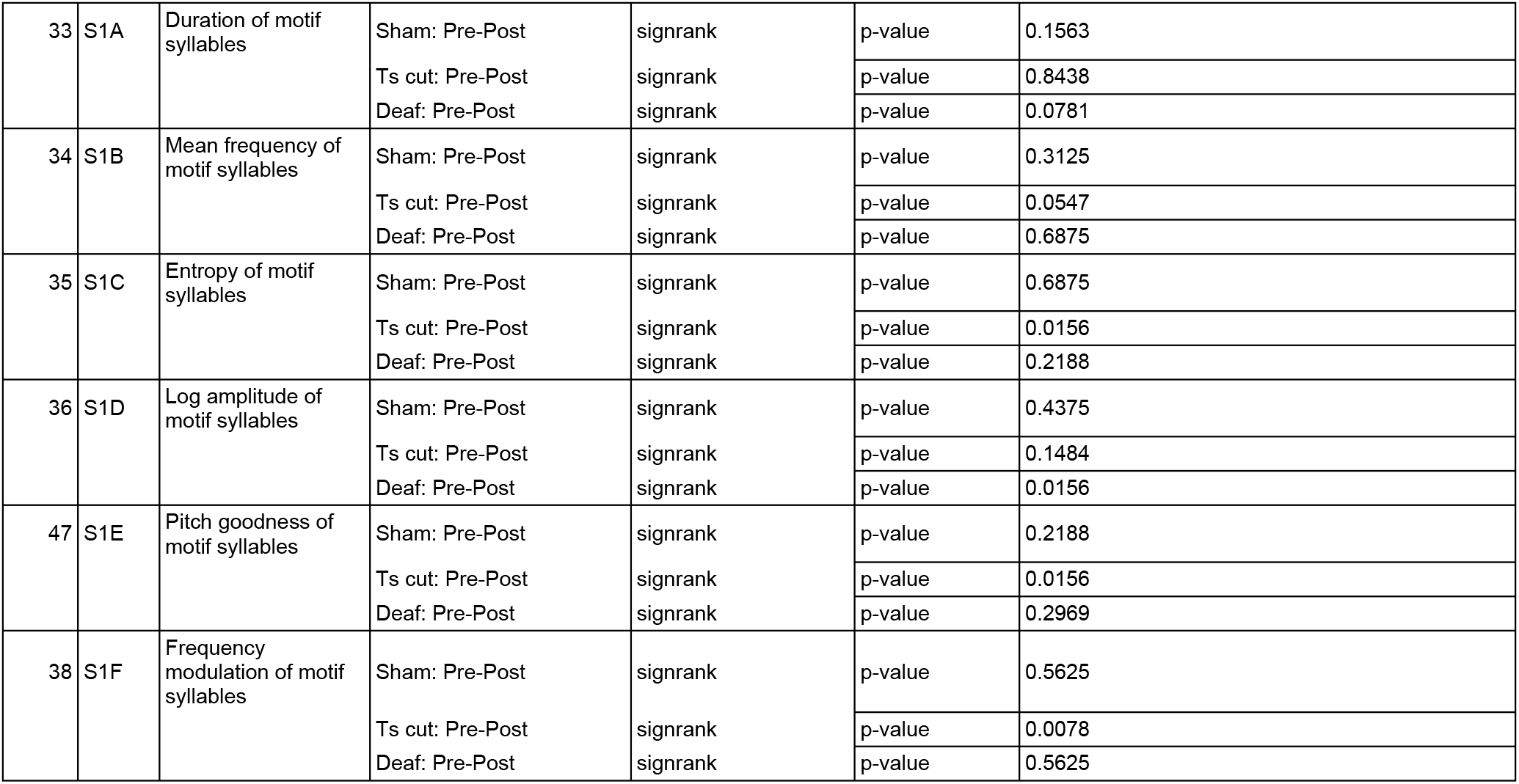
Details of statistical tests and the associated p-values for all analyses

